# Chromatin 3D interaction analysis of the *STARD10* locus unveils *FCHSD2* as a new regulator of insulin secretion

**DOI:** 10.1101/2020.03.31.017707

**Authors:** Ming Hu, Inês Cebola, Gaelle Carrat, Shuying Jiang, Sameena Nawaz, Amna Khamis, Mickael Canouil, Philippe Froguel, Anke Schulte, Michele Solimena, Mark Ibberson, Piero Marchetti, Fabian L. Cardenas-Diaz, Paul J. Gadue, Benoit Hastoy, Leonardo Alemeida-Souza, Harvey McMahon, Guy A. Rutter

**Affiliations:** Section of Cell Biology and Functional Genomics, Department of Medicine, Imperial College London, Hammersmith Hospital, Du Cane Road, London W12 0NN, United Kingdom; Section of Genetics and Genomics, Department of Metabolism, Digestion and Reproduction, Imperial College London, Hammersmith Hospital, Du Cane Road, London W12 0NN, United Kingdom; Oxford Centre for Diabetes Endocrinology and Metabolism, University of Oxford, Churchill Hospital, Headington, Oxford OX3 7LE Oxford; Univ. Lille, CNRS, CHU Lille, Institut Pasteur de Lille, UMR 8199 - EGID, F-59000 Lille, France; Sanofi-Aventis Deutschland GmbH, 65926 Frankfurt am Main, Germany; Paul Langerhans Institute of the Helmholtz Center Munich at the University Hospital and Faculty of Medicine, TU Dresden, 01307 Dresden, Germany; Vital-IT Group, SIB Swiss Institute of Bioinformatics, 1015 Lausanne, Switzerland; Department of Endocrinology and Metabolism, University of Pisa, 56126 Pisa, Italy; Department of Pathology and Laboratory Medicine, University of Pennsylvania, Philadelphia; and Centre for Cellular and Molecular Therapeutics, Children’s Hospital of Philadelphia, PA. United States of America; HiLIFE Institute of Biotechnology & Faculty of Biological and Environmental Sciences, University of Helsinki, Helsinki, Finland; MRC MRC Laboratory of Molecular Biology, Francis Crick Avenue, Cambridge, UK CB2 0QH; Lee Kong Chian School of Medicine, Nan Yang Technological University, Singapore

**Author notes:** Correspondence to Professor Guy A. Rutter.

**Keywords:** GWAS, genetic variant, enhancer cluster, chromatin structure, gene regulation, *STARD10*, *FCHSD2*, insulin secretion

## Abstract

Genome-wide association studies have identified thousands of genetic variants associated with type 2 diabetes (T2D) risk. Using chromatin conformation capture we show that an enhancer cluster in the *STARD10* T2D locus forms a defined 3D chromatin domain. A 4.1 Kb region within this region, carrying five disease-associated variants, physically interacts with CTCF-binding regions and with an enhancer possessing strong transcriptional activity. Analysis of human islet 3D chromatin interaction maps identifies *FCHSD2* gene as an additional target of the enhancer cluster. CRISPR-Cas9-mediated deletion of the variant region, or of an associated enhancer, in human pancreatic beta cells impaired glucose-stimulated insulin secretion. Expression of both *STARD10* and *FCHSD2*, but not *ARAP1*, was reduced in cells harboring CRISPR deletions, and expression of *STARD10* and *FCHSD2* was associated with the possession of variant alleles in human islets. Finally, CRISPR-Cas9-mediated loss of *STARD10* or *FCHSD2* impaired regulated insulin secretion. Thus, multiple genes at the *STARD10* locus influence β cell function.

## INTRODUCTION

Genome-wide association studies (GWAS) have identified over 400 genetic signals across >200 loci that associate with type 2 diabetes (T2D) risk (Morris et al., 2012; DIAbetes Genetics Replication And Meta-analysis (DIAGRAM) Consortium et al., 2014; Voight et al., 2010; Scott et al., 2017 and Mahajan et al., 2018). Data from these and other studies indicate that islet dysfunction plays a major role in T2D genetic risk. However, associated genetic variants usually lie in intergenic or intronic regions of the genome, and only a minority affect protein sequences (Fuchsberger et al., 2016).

One plausible mechanism by which genetic variants may contribute to T2D risk is by affecting functional noncoding sequences. Consisting of short DNA regions, and located at varying distances from promoter sequences, enhancers are *cis*-regulatory elements that promote the expression of target genes due to their co-occupancy by tissue-enriched transcription factors and coactivators. T2D GWAS variants are enriched within pancreatic islet enhancer clusters, also termed “clusters of open regulatory elements” (COREs), stretch enhancers, super-enhancers and, more recently, enhancer hubs (Gaulton et al., 2010; Parker et al., 2013; Pasquali et al., 2014 and (Miguel-Escalada et al., 2019). Enhancer clusters often control temporal and cell-type specific functions and define cell identity (Gosselin et al., 2014; Huang et al., 2016 and Whyte et al., 2013). Thus, genetic variants in islet enhancer clusters may contribute to diabetes risk by perturbing islet transcriptional networks. Consequently, in addition to the identification of causal variants, functional characterization of enhancer-target gene(s) interactions, and of their effect(s) on β cell function, are required to fully understand the genetic influence of T2D pathogenesis.

Enhancers interact with target gene(s) to regulate their expression, an effect achieved through chromatin looping, often mediated by the highly conserved architectural protein CTCF (CCCTC-binding factor) (Bonev and Cavalli, 2016 and Williams and Flavell, 2008). CTCF contains a DNA-binding domain that recognizes a non-palindromic motif. Highlighting the relevance of CTCF sites in chromatin architecture and gene regulation, deletion or inversion of CTCF binding sites can affect chromatin looping and cause dysregulated gene expression (de Wit et al., 2015; Guo et al., 2015 and Mandegar et al., 2016).

Several GWAS variants, including rs7903146 in the *TCF7L2* gene, are enriched in regions of open chromatin and confer allele-specific activity to an intronic islet enhancer (Gaulton et al., 2010). Similarly, a T2D-associated risk variant, rs7163757, in the *VPS13C/C2CD4A*/*C2CD4B* locus on chromosome 15 (Strawbridge et al., 2011) resides in an islet specific enhancer cluster and affects the expression of nearby genes (Mehta et al., 2016 and Kycia et al., 2018).

In our recent studies (Carrat et al., 2017), we used functional genome-wide association (fGWAS) (Pickrell, 2014) to fine map a diabetes-associated credible set in the *ARAP1/STARD10* T2D GWAS locus, in which the risk haplotype has a global frequency of 86%. The identified credible set is composed of eight variants, of which five displayed a posterior probability >0.05, in intron 2 of the *STARD10* gene. One of these (Indel rs140130268), which possessed the highest probability, is located at the edge of a region of open chromatin (ATAC-seq). Whether, and how, these variants affect the expression of local or remotely-located genes in human β cells was not, however, examined in our earlier report.

In the present study, we have used human EndoC-βH1 cells, which recapitulate many of the functional properties of native human β cells (Ravassard et al., 2011), and deployed chromatin interaction analyses and β-cell tailored CRISPR-Cas9 genome editing to explore this question. We show that the variant region is required for normal glucose-stimulated insulin secretion and identify the enhancer regions with which it interacts physically. We also demonstrate direct roles for *STARD10* in human-derived β cell function. Finally, we provide genetic and functional evidence of a role for a previously unimplicated nearby gene, *FCHSD2*, encoding a regulator of membrane trafficking and endocytosis (Almeida-Souza et al., 2018) in variant action.

## RESULTS

### Chromatin landscape at the *STARD10* locus

We investigated regulatory regions at the T2D GWAS locus close to *STARD10* by overlaying multiple human islet epigenomic datasets: Assay for Transposase-Accessible Chromatin (ATAC-seq), histone marks associated with active chromatin (i.e. H3K27ac) and ChIP-seq for key islet transcription factors (TFs) (Pasquali et al., 2014 and Miguel-Escalada et al., 2019). This analysis revealed multiple regulatory elements (R1-13) that are active in human islets, including a cluster of six enhancers (Figure 1A). Several of these were bound by islet-enriched TFs such as NKX2.2, FOXA2 and MAFB, and are thus likely to contribute to an islet-specific gene expression signature. We also detected two binding sites for the chromatin architectural factor CTCF flanking the enhancer cluster, which may be involved in the creation of a distinct chromatin domain and mediate long-range looping with distal target genes (Figure 1A).

**Figure 1.**
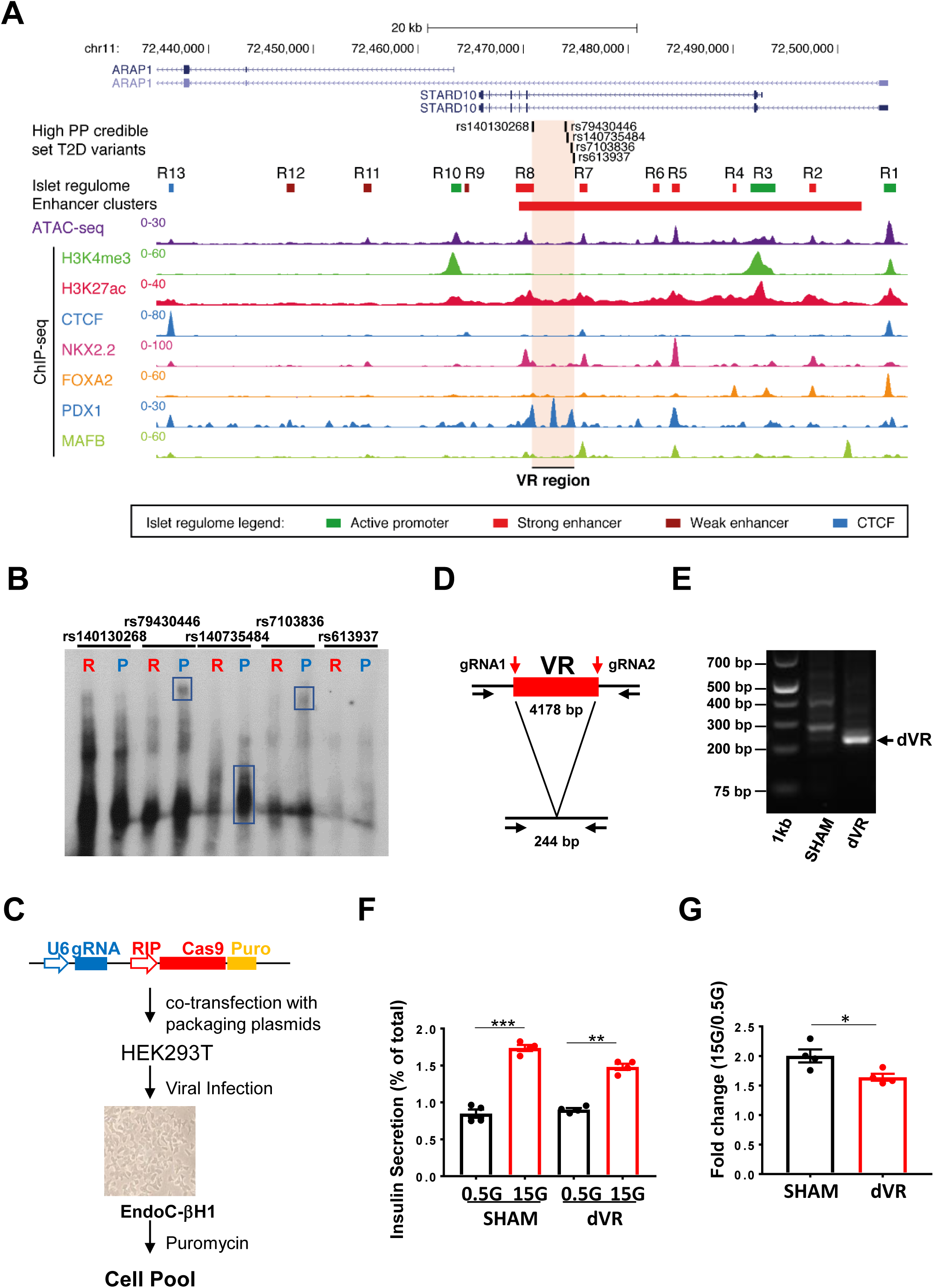
Variant region (VR) in local chromatin structure and β cell function. (A) Epigenomic map of *STARD10* locus in human islets. The open chromatin regions identified by ATAC-seq were termed and labelled as R for regulatory region. Enhancer cluster: solid red bar; Variant region: VR. Active enhancer regions: R8, R7, R6, R5, R4 and R2. (B) Electrophoretic Mobility Shift Assay (EMSA). R: risk allele; P: protective allele. Data are from two independent experiments. (C) Diagram of CRISPR-Cas9 mediated genome editing with a β cell-tailored vector via lentiviral approach. Lentiviruses were generated in HEK293T cells, titrated and used to infect EndoC-βH1 cells. Puromycin was used to select viral resistant cells and generate a cell pool. (D) Strategy of variant region (VR) deletion by CRISPR-Cas9 genome editing. Two gRNAs were designed to flank the VR region and generate a 4178 bp deletion. (E) Electrophoresis of PCR products amplified from SHAM and VR-deleted (dVR) genomic DNA. Note, the bands in the SHAM lane (around 300-400 bp) were non-specific products. (F) Representative data of glucose stimulated insulin secretion (GSIS) assay. Cells were cultured at basal level glucose (0.5 mM) condition and then stimulated with high glucose (15 mM). Data are mean ± SEM. *, *P* < 0.05; **, *P* < 0.01; ***, *P* < 0.005. *n* = 4. (G) Fold change of secreted insulin. Data are normalized to insulin secretion at basal level (0.5G). *, *P* < 0.05; **, *P* < 0.01; ***, *P* < 0.005. *n* = 4.

### Credible set variants exhibit differential transcription factor binding and transcriptional activity

We next turned our attention to the five variants in the credible set with greatest causal probability, as defined previously by fine mapping and fGWAS analysis (Carrat et al., 2017). These variants span a 4.1 Kb interval and include two deletions (indels: rs140130268 and rs140735484) and three single nucleotide polymorphisms (SNP; rs79430446, rs7103836 and rs613937). Of these, indel rs140130268 displayed the highest posterior probability and showed allele-specific transcriptional activity in β cells (Carrat et al., 2017). Detailed epigenomic analysis (Figure 1A) mapped these variants to the center of an enhancer cluster defined by strong H3K27ac enrichment in islet chromatin (Pasquali et al., 2014), though none of them resided within a previously mapped open chromatin region. We note, however, that the risk haplotype in this particular locus, which associates with lowered regulatory activity (Carrat et al., 2017), is present in 86 % of the human population. Thus, it is possible that the existing regulatory maps do not represent the epigenomic landscape of carriers of the non-risk haplotype (MAF = 0.14).

A likely mechanism by which the risk haplotype could confer reduced local chromatin accessibility is via the alteration of transcription factor recognition sequences. To test this hypothesis, we performed TF motif analysis (Khan et al., 2018) on these variants, which suggested that four of the five variants may affect TF binding to this enhancer cluster (Figure S1 and Table S2).

To further explore this possibility, we assayed allele-specific transcription factor binding by Electrophoretic Mobility Shift Assays (EMSA) with oligonucleotides carrying either risk or protective variants. While differences were modest between risk and protective alleles for rs140130268, we observed marked differences in DNA-protein complex mobility between risk and protective alleles for rs79430446, rs140735484 and rs7103836 (Figure 1B). These results therefore point to a potential regulatory function of several of the variants in this credible set.

### Deletion of the variant region from the EndoC-βH1 genome reduces insulin secretion

To assess the collective importance of the above variants, we deleted the entire 4.1 Kb variant region in EndoC-βH1 cells, which are homozygous for the risk haplotype, using CRISPR-Cas9-mediated genome editing (Figure 1C). To this end, we designed two gRNAs flanking the 4.1 kb genomic region that contains the T2D variants (Figure 1D). CRISPR-Cas9 genome editing was then performed in the glucose-responsive human β cell line EndoC-βH1 (Ravassard et al., 2011), delivering the guide RNAs and the Cas9 gene under the control of the rat insulin promoter (RIP), and a puromycin resistance cassette. Control (SHAM) cells were treated with lentivirus lacking gRNAs and selected with puromycin in the same way. Deletion of the variant region was confirmed by genomic PCR (Figure 1E) and Sanger sequencing (Figure S1B). Quantification by qPCR revealed the remaining wildtype allele in dVR cells represented ∼ 48 % (48.04 ± 2.17) of total. In addition, we also detected DNA inversion after editing, corresponding to ∼ 10 % (9.5 ± 0.27) of the remaining wildtype alleles. Hence, the overall deletion efficiency was about 47 % (Figure S1C and S1D).

To determine whether loss of the T2D variants may impact β cell function, we assayed glucose-stimulated insulin secretion (GSIS) in the presence (SHAM) or absence (dVR) of the variant region. dVR cells displayed a small but significant reduction in GSIS (fold change: SHAM: 2.00 ± 0.11 vs. dVR: 1.64 ± 0.06. P = 0.0267, *n* = 4) (Figure 1F and 1G).

### The T2D credible set variants interact with active islet regulatory elements

The 4.1 kb region (VR) that encompasses the five T2D credible set variants lies between two active enhancers (R8 and R7, Figure 1A), but does not overlap with any annotated islet regulatory element. Since the above experiments demonstrated that this region is involved in the regulation of insulin secretion (Figure 1F and 1G) we hypothesized that it may contribute to β cell function by affecting chromatin topology and/or gene expression. To determine which genomic region(s) the VR may interact with, we performed circular chromosome conformation capture (4C) analysis (Göndör et al., 2008) (Figure 2A). Out of a total of 56 clones obtained after restriction enzyme digestion (*Pst*I and *Msp*I), and DNA ligations, we detected four clones containing a DNA fragment within region R13, which corresponds to a strong CTCF site in human islets (Figure 2B). Moreover, five sequenced fragments mapped to a region 1.3 kb upstream of the R1 region, which contains one of the two promoters of *STARD10* that are active in human islets, and is bound by CTCF (Figure 2B). The remaining fragments corresponded to DNA regions in the vicinity of the viewpoint, likely reflecting local chromatin collisions (Hagège et al., 2007).

**Figure 2.**
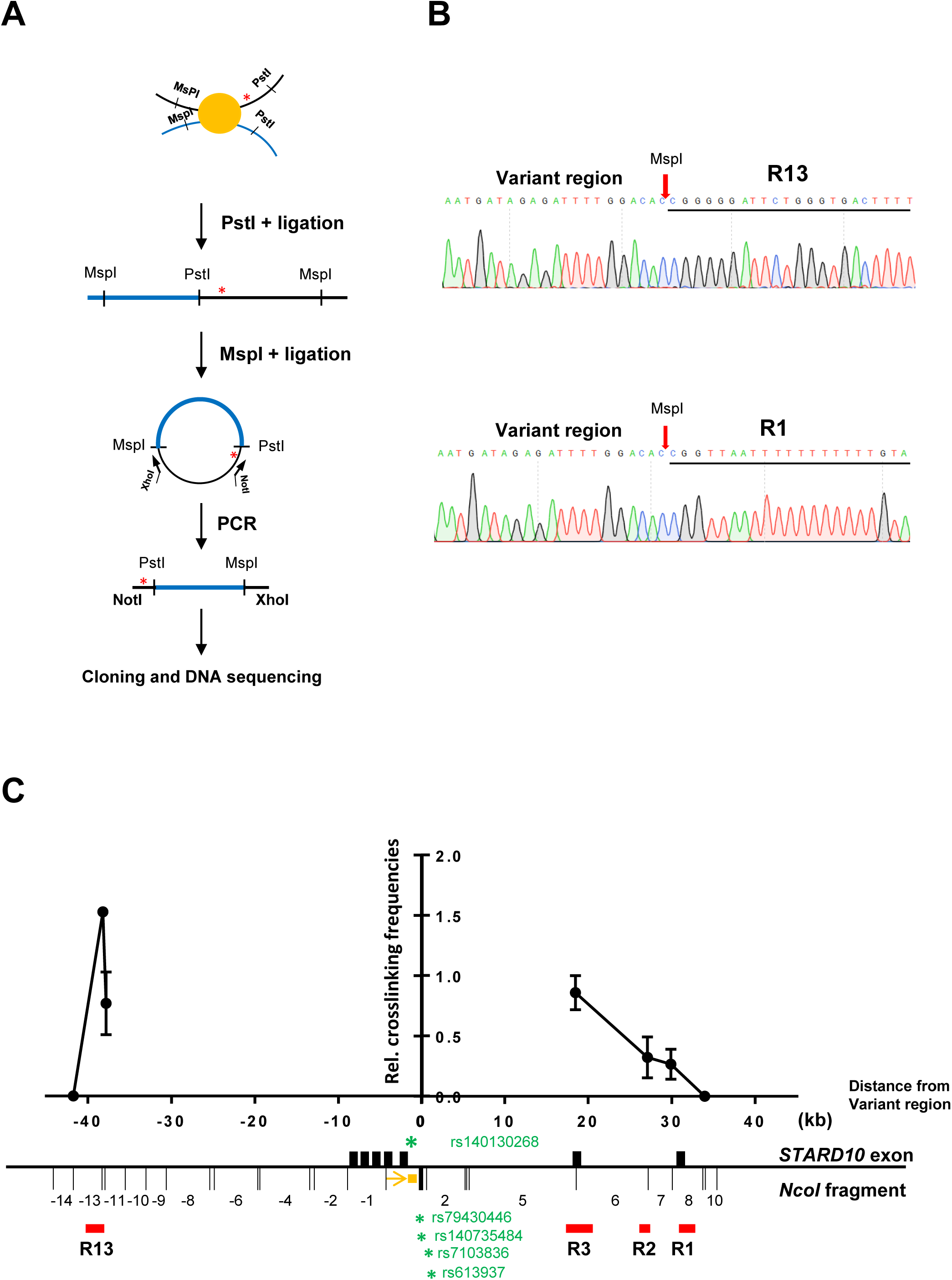
Identification and confirmation of genomic regions that physically interact with variant region. (A) Diagram of conventional 4C assay. (B) Representative DNA sequencing results showing ligation products between variant region and interactive regions (R13 and R1 regions). (C) Representative 3C-qPCR data for chromatin interactions at *STARD10* locus. Viewpoint: VR region; Black bar: *STARD10* exons; red bar: regulatory regions; orange box: qPCR probe; orange arrow: qPCR constant primer; green stars: credible set genetic variants. The numbering of *NcoI* fragments is given relative to the viewpoint. The leftmost dot corresponds to the 5’ end of -13 DNA fragment. Note, the bait fragment contains rs140130268 only. rs79430446 and rs140735484 are in *Nco*I fragment 1; rs7103836 and rs613937 are in *NcoI* fragment 2. R13 is in *NcoI* fragment -13 and -12, while R1 is in *NcoI* fragment +8. Data were normalized to a *CXCL12* loading control. *n* = 3.

To validate the 4C results above, we performed 3C analysis (Figure 2C). Taking the T2D credible set variants region as a viewpoint, we detected higher interaction frequencies of the variant region with both the CTCF site R13 and the two promoters of *STARD10* (R3 and R1). Taken together, these results demonstrate that the T2D credible set variants region in *STARD10* undergo cis-interactions with human islet regulatory elements, including CTCF anchor points and the two promoters of *STARD10*.

### Identification of CTCF binding sites (CBSs) at the *STARD10* locus

Inspection of human islet ChIP-seq datasets, together with *in silico* transcription factor binding motif analysis (Figure 1A and Figure S3A), revealed that both R1 and R13 contain binding sites for critical islet transcription factors, including NKX2.2 and FOXA2, indicating their potential role in the regulation of islet gene expression. Of note, the two regions also showed enrichment for CTCF by ChIP-seq in human islets (Figure 1A). Using ChIP-qPCR assay in EndoC-βH1 cells, we detected binding of CTCF to five out of the eight potential CTCF binding motifs (Figure 3A and 3B). Importantly, we detected CTCF enrichment at convergent CTCF binding motif sequences, a configuration that has been previously shown to be involved in chromatin looping (de Wit et al., 2015).

**Figure 3.**
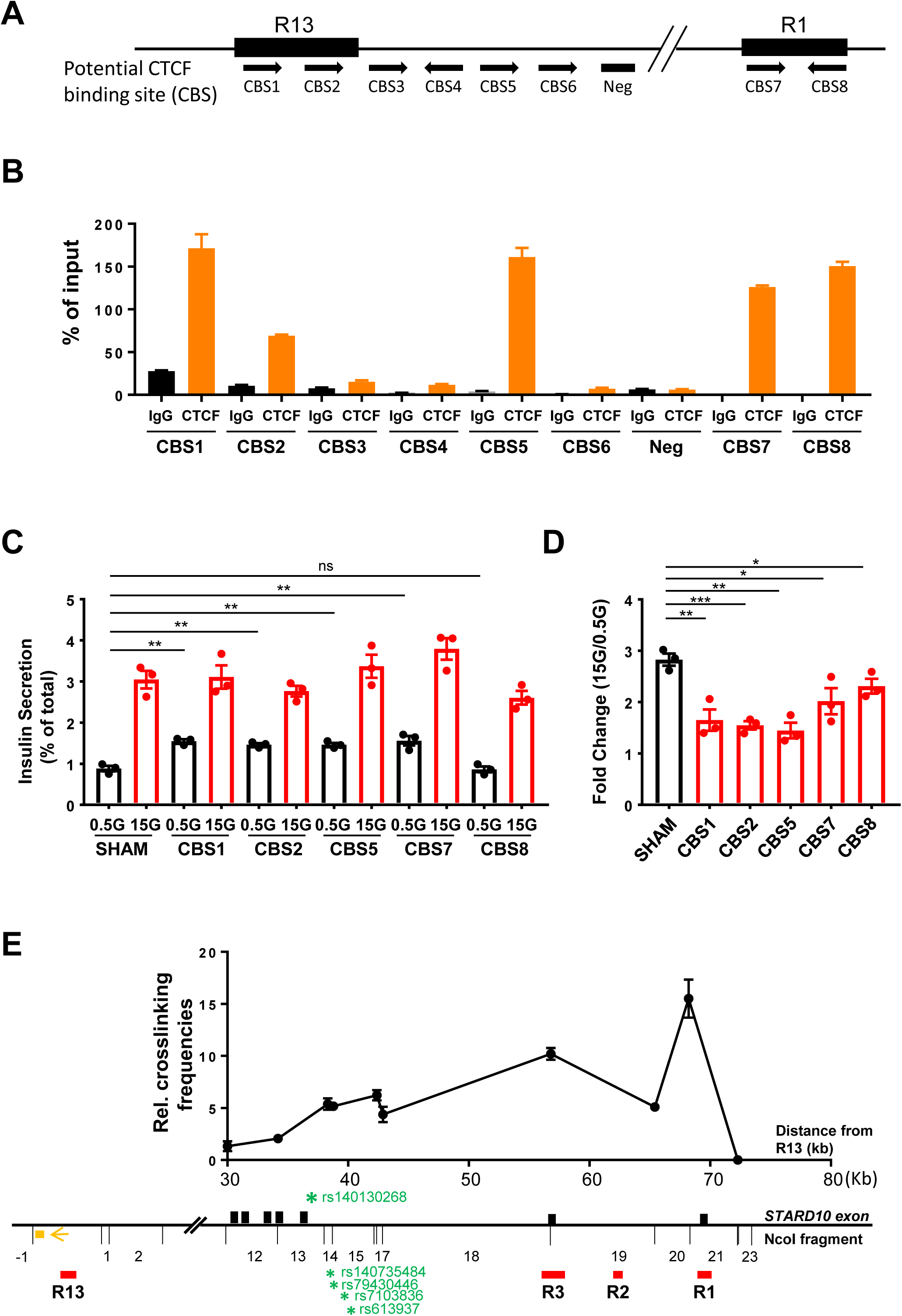
CTCF binding sites (CBSs) at R13 and R1 regions. (A) Diagram of potential CTCF binding site (CBS) within and surrounding R13 and R1 regions. Black arrows: binding orientation of CTCF. (B) Representative data of ChIP-qPCR analysis for CTCF binding at predicted binding sites. *, *P* < 0.05; **, *P* < 0.01; ***, *P* < 0.005. *n* = 3. (C) Representative data of GSIS assay. Data are mean ± SEM. *, *P* < 0.05; **, *P* < 0.01; ***, *P* < 0.005. *n* = 3. (D) Fold change of secreted insulin. Data are normalized to insulin secretion at basal level (0.5 mM glucose). Data are mean ± SEM. *, *P* < 0.05; **, *P* < 0.01; ***, *P* < 0.005. *n* = 3. (E) Representative 3C-qPCR data of chromatin interactions at *STARD10* locus. The numbering of *NcoI* DNA fragments is given relative to the viewpoint. Viewpoint: R13 region; Black bars: *STARD10* exon; red bars: regulatory region; orange box: qPCR probe; orange arrow: qPCR constant primer; green stars: credible set genetic variants. Note, R1 is in the *NcoI* fragment +21. Data were normalized to a *CXCL12* loading control. *n* = 3.

### Mutation of CBSs leads to impaired insulin secretion

Higher order chromatin structure is required for the regulation of cell-specific transcriptional activity (de Wit et al., 2015; Guo et al., 2015 and Lupiáñez et al., 2015). Given the structural features of the *STARD10* locus described above, we investigated whether loss of key architectural elements in the *STARD10* locus could lead to β cell function impairment. Using CRISPR-Cas9, we mutated the identified CBSs in the *STARD10* locus individually in EndoC-βH1 cells. ChIP-qPCR for CTCF confirmed significant loss of CTCF binding ability at designated binding sites after CRISPR targeting (Figure S3B). In assays of glucose-stimulated insulin secretion (GSIS), we observed that CRISPR-Cas9-mediated targeting of four out of the five CBSs in the region (CBS1, CBS2, CBS5 and CBS7) led to increased basal insulin secretion (at 0.5 mM glucose) and lowered the fold change in secretion at high (0.5 vs. 15 mM) glucose (Figure 3C and 3D). Similar results were observed for insulin secretion stimulated by cyclic AMP (cAMP)-raising reagents such as isobutylmethylxanthine (IBMX) and forskolin, as well as in response to cell depolarization with KCl (Figure S3C and S3D). Furthermore, we found that the expression patterns of *STARD10* and of nearby genes was significantly altered in CBS mutated cells. qRT-PCR analyses (Figure S3E) revealed that *STARD10, ATG16L2* and *FCHSD2* were the most downregulated in CBS mutant cells while *ARAP1*, a gene that resides near the T2D-associated credible set was unaffected.

Taken together, these results demonstrate that CTCF binding, through its likely impact on chromatin structure organization at the *STARD10* locus, is necessary to maintain normal β cell function.

### R13 and R1 regions form chromatin loop via CTCF binding sites

CTCF, together with the Cohesin complex, plays an important role in the formation of higher order chromatin structures and may act as an insulator or boundary between cis-regulatory elements and their target genes (Bell et al., 1999; Lupiáñez et al., 2015 and Merkenschlager and Odom, 2013). As shown in Figure 2C, the region containing the T2D credible set in *STARD10* interacts with both R1 and R13 (and both of the latter contain *bona fide* CTCF convergent binding sites) (Figure 3A). We therefore hypothesized that the two regions may interact with each other, via the formation of a CTCF-CTCF loop, to establish a restricted chromatin domain. To test this hypothesis, we performed 3C analysis in EndoC-βH1 cells and explored the interaction frequency between R1 and R13. Taking R13 as the viewpoint, we detected interaction frequencies above background level across the entire *STARD10* locus (Figure 3E), particularly with the two promoters of *STARD10* (R3 and R1), but also with the T2D credible set region, as observed previously (Figure 2C). Together, these observations confirm that the T2D variant region interacts with a distal CTCF site in β cells, and demonstrate that R13 and R1 are also associated through chromatin looping.

### Screen of annotated genomic features reveals a functional islet enhancer that regulates basal insulin secretion

Since the T2D variants are deeply embedded within the region of active enhancers (Figure 1A), we hypothesized that the causal variants may exert their effect(s) by altering the activities of these enhancers. To explore this possibility, we sought first to understand the roles of the enhancer cluster in the control of β cell gene expression and function.

Regulome analyses of human islet samples, including ATAC-seq and ChIP-seq for H3K27ac, revealed six active enhancers showing islet TF binding (Figure 1A). We thus tested these regions by luciferase reporter assay in EndoC-βH1 cells, which revealed that R2, a small enhancer located between the two *STARD10* islet promoters, had 6.25-fold activity increase compared with control (Control: 1.26 ± 0.06 vs. R2: 7.82 ± 0.17. P < 0.0001, *n* = 3) (Figure 4A). Other putative enhancers displayed negligible activity in this assay.

**Figure 4.**
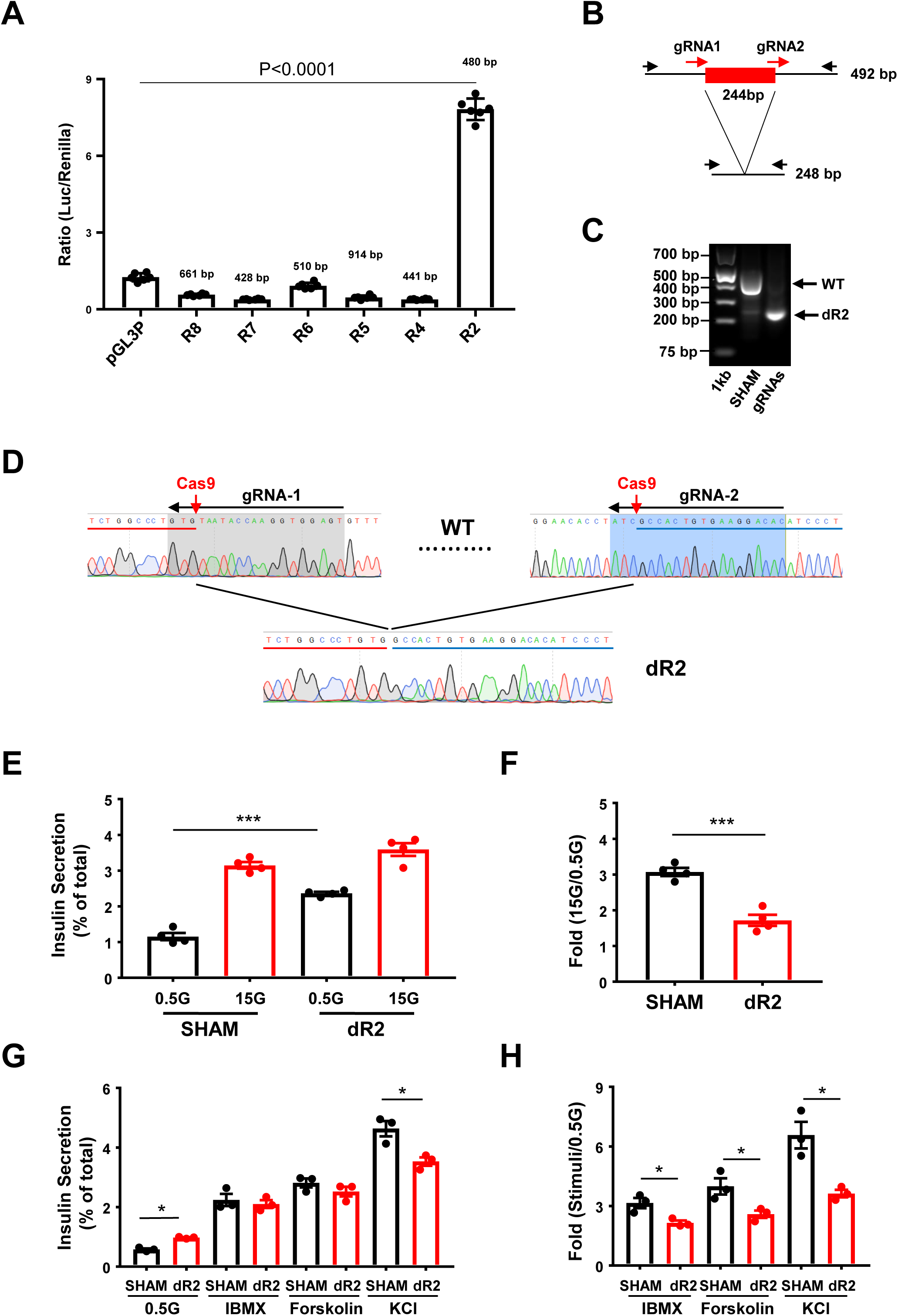
Role of enhancer R2 in insulin secretion. (A) Promoter-luciferase assay in EndoC-βH1 cells. ***, *P* < 0.0001. *n* = 3. (B) Strategy of R2 deletion by CRISPR-Cas9 genome editing. Two gRNAs were designed to delete 244 bp of core R2 region. (C) Gel electrophoresis of PCR products amplified from genomic DNAs isolated from control and R2-deleted (dR2) cells. (C) Sanger sequencing confirmation of R2 deletion. Red and blue bars represent the 5’ and 3’ end of DNA sequence flanking R2 region. (E) Representative data of GSIS assay. Data are mean ± SEM. *, *P* < 0.05; **, *P* < 0.01; ***, *P* < 0.005. *n* = 4. (F) Fold change of secreted insulin. Data are normalized to insulin secretion at basal level (0.5 mM glucose). *, *P* < 0.05; **, *P* < 0.01; ***, *P* < 0.005. *n* = 4. (G) Representative data of insulin secretion stimulated by other stimuli. Data are mean ± SEM. *, *P* < 0.05; **, *P* < 0.01; ***, *P* < 0.005. *n* = 3. (H) Fold change of secreted insulin. Data are normalized to insulin secretion at basal level (0.5 mM glucose). *, *P* < 0.05; **, *P* < 0.01; ***, *P* < 0.005. *n* = 3.

The R2 enhancer contains several recognition sequences for the binding of islet-associated TFs, such as FOXA2 and PAX4, suggesting a critical role in the regulation of nearby β cell genes (Figure S4A). To assess the role of this enhancer in β cell function, we deleted the core region of R2 (244 bp) from EndoC-βH1 cells using CRISPR-Cas9 (Figure 4B), achieving ∼85 % loss of targeted alleles (85.67 ± 1.44) (Figure 4C, 4D and Figure S4B, S4C, S4D). GSIS was significantly impaired in edited versus sham-treated cells (Fold change [15 mM glucose *vs* 0.5 mM glucose]: SHAM: 3.07 ± 0.11 vs. dR2: 1.72 ± 0.16. P = 0.0004, *n* = 4) (Figure 4E and 4F), largely due to increased basal insulin secretion (0.5 mM glucose) [SHAM: 1.16 ± 0.10 vs. dR2: 2.36 ± 0.04. P = 0.0004, *n* = 4]. The stimulation of insulin secretion was even more sharply reduced in R2-deleted cells in response to cAMP-raising agents or after depolarization with KCl to prompt Ca^2+^ entry (Figure 5G and 5H).

**Figure 5.**
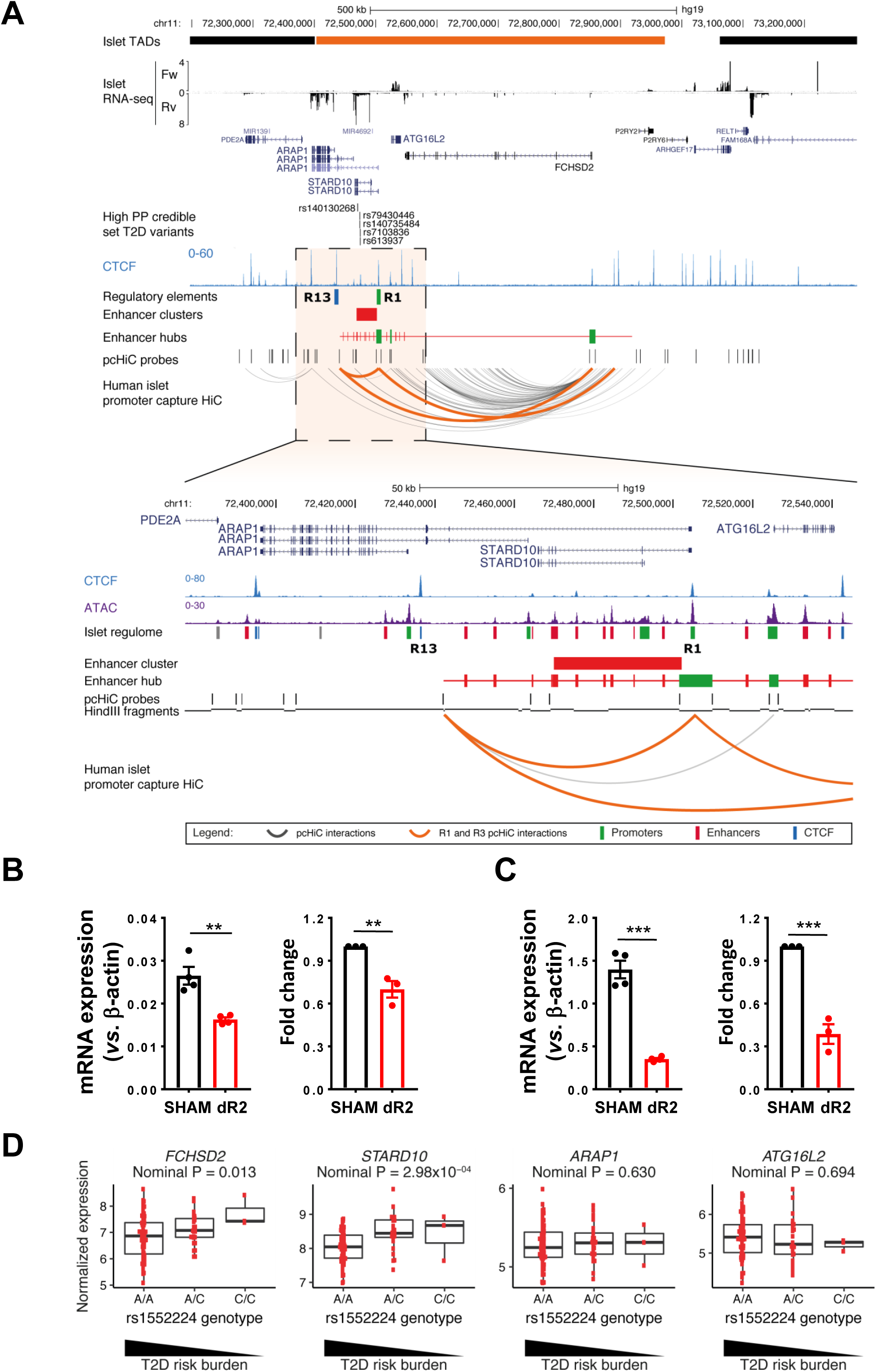
Target genes regulated by the enhancer cluster. (A) Human islet promoter capture-HiC (pcHi-C) map at *STARD10* locus and surrounding region. pcHi-C interactions were considered significant with CHiCAGO > 5. Orange interactions in the pcHiC track depict interactions mediated by the CTCF sites that flank the enhancer cluster (R1 and R13). All other significant interactions are shown in dark grey. Enhancer hubs track shows all enhancers (red) and promoters (green) contained within the *STARD10* hub.Orange bar in islet TADs track highlights the TAD encompassing *STARD10* and surrounding genes (*ARAP1, ATG16L2*, and *FCHSD2*). (B and C) Taqman qRT-PCR analysis of gene expression in sham control and R2-deleted (dR2) cells. *, *P* < 0.05; **, *P* < 0.01; ***, *P* < 0.005. *n* = 3. (D) eQTL analysis of human islet samples. Y-axis represents normalized intensities (using RMA method) from the Affymetrix Human Genome U133 Plus 2.0 Array. Total human samples is 203.

### Enhancer cluster regulates *FCHSD2* gene expression through chromatin looping

Individual enhancers can regulate multiple genes within the same cellular population, forming distinct 3D chromatin regulatory domains (“enhancer hubs”) (Miguel-Escalada et al., 2019) and Oudelaar et al., 2018). We therefore postulated that the *STARD10* enhancer cluster might be part of a broader 3D chromatin domain in human islets.

To assess this possibility, we queried the recently published genome-wide map of human islet 3D chromatin interactions (promoter-capture HiC, pcHi-C) (Miguel-Escalada et al., 2019). This analysis revealed that *STARD10* resides within an islet enhancer hub, together with *ARAP1, ATG16L2* and the distal gene *FCHSD2*, located ∼400 kb downstream of the *STARD10* enhancer cluster (Figure 5A). These four hub genes are expressed at relatively high to medium levels in human islets: *ARAP1, STARD10, ATG16L2* and *FCHSD2* (see RNA-seq track in Figure 5A, and Figure S5C). We note, however, that ARAP1 has three annotated promoters and only one of them is included in the hub (Miguel-Escalada et al., 2019) (Figure S5C). This promoter drives the expression of a longer isoform of *ARAP1* and is shared with *STARD10.* Examination of human islet RNA-seq revealed that the longer *ARAP1* transcript isoform displays much lower expression in human islets than the two other isoforms (Figure S5C), possibly reflecting CTCF binding at this site. Thus, most of the *ARAP1* transcription is driven by promoters P2 and P3 (Figure S5C), which reside outside of the *STARD10* islet enhancer hub.

To identify the gene(s) that are regulated by the nearby enhancer cluster, we carried out gene expression profiling in R2-deleted (dR2) cells. This revealed significant down-regulation of only *FCHSD2* (Fold change: SHAM: 1.0 ± 0 vs. dR2: 0.7 ± 0.06, P = 0.0066, *n* = 3) and *STARD10* (Fold change: SHAM: 1.0 ± 0 vs. dR2: 0.39 ± 0.07. P = 0.0008, *n* = 3) (Figure 5B and 5C). In contrast, *ATG16L2* and *ARAP1*, the gene which was closest in linear distance to the enhancer cluster, were not affected by R2 deletion (Figure S5A and S5B). The latter observation is in line with our previous analysis of islet expression quantitative trait loci (eQTL), which did not reveal any association between the *STARD10* T2D variants in this locus and the expression of *ARAP1* (Carrat et al., 2017). Further analysis of the pcHi-C dataset confirmed that regions R1 and R13 interact in human islet chromatin (Figure 5A), consistent with our 3C analysis in EndoC-βH1 cells (Figure 3E). Moreover, the pcHi-C dataset revealed that the two CTCF binding sites that flank the *STARD10* enhancer cluster (R1 and R13) undergo long-range interactions with the promoter region of *FCHSD2* (Figure 5A).

### Human islet expression quantitative trait loci (eQTL)

To gain insight into the relevance of our findings in the context of human islet physiology and diabetes risk, we analyzed the expression of *FCHSD2* in a cohort of 103 subjects who provided pancreatic samples after partial pancreatectomy and laser capture microdissection analysis (IMIDIA consortium; 47 non-diabetic, 56 T2D) (Solimena et al., 2018 and Marchetti et al., 2018). We observed lower *FCHSD2* expression in carriers of the risk alleles of variants (rs75896506, rs11603334, rs1552224, Nominal P-value = 0.013), which are in high LD with the high posterior probability variant rs140130268 (EUR R2 = 0.8934) (Figure 5D and Table S1). Consistent with earlier reports (Carrat et al., 2017 and Miguel-Escalada et al., 2019), the risk variants also associated with lower *STARD10* levels in this cohort (Nominal P-value = 2.98 × 10^−4^) (Figure 5D and Table S1). In contrast, no eQTL was found for *FCSHD2* in organ donor (OD) -obtained samples from the same cohort (p=0.89), whilst the signal for *STARD10* remained nominally significant (p = 2.19 × 10^−3^). These results suggest that the T2D-associated variants in the *STARD10* enhancer hub selectively associate with differential expression of *STARD10* and *FCHSD2*.

### *STARD10* and *FCHSD2* knockout affect regulated insulin secretion in human β cells

We have previously demonstrated the importance of the *STARD10* gene in regulating insulin processing and secretion in the mouse (Carrat et al., 2017). The findings above suggest that FCHSD2 may also play a role in the control of insulin secretion. To further explore the potential roles of *STARD10* and *FCHSD2* in controlling insulin secretion in the human setting, we deployed CRISPR-Cas9 genome editing in EndoC-βH1 cells to generate frameshift mutations in exon 2 of *STARD10* (Figure 6A and Figure S6A) and exon 1 of *FCHSD2* (Figure 6E and Figure S6B). Western blot analysis confirmed the expected loss of protein expression with 80-90 % efficiency for both STARD10 (Figure 6B) and FCHSD2 (Figure 6F).

**Figure 6.**
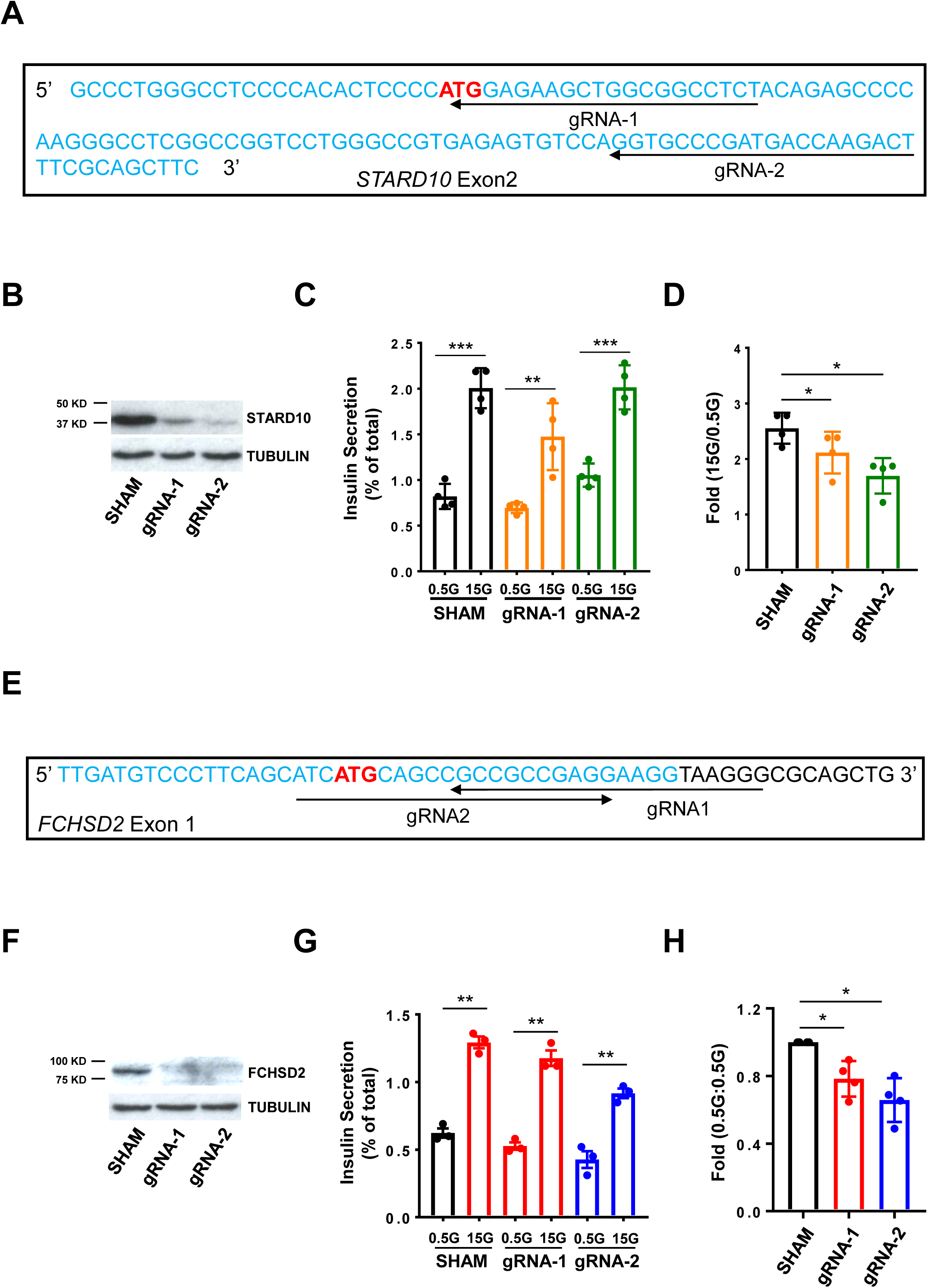
Effect of *STARD10* and *FCHSD2* gene knockout on insulin secretion. (A) gRNA design at *STARD10* exon2 region. ATG: start codon. gRNA1 and gRNA2 were delivered individually into EndoC-βH1 cells. (B) Western blot analysis of STARD10 protein in *STARD10*-KO cells. Tubulin was served as a protein loading control. (C) Representative data of GSIS assay in *STARD10*-KO cells. Data are mean ± SEM. *, *P* < 0.05; **, *P* < 0.01; ***, *P* < 0.005. *n* = 4. (D) Fold change of secreted insulin. Data are normalized to insulin secretion at basal level (0.5G). *, *P* < 0.05; **, *P* < 0.01; ***, *P* < 0.005. *n* = 4. (E) gRNA design at *FCHSD2* exon1 region. ATG: start codon. gRNA1 and gRNA2 were delivered individually into EndoC-βH1 cells. (F) Western blot analysis of FCHSD2 protein in *FCHSD2*-KO cells. Tubulin was served as a protein loading control. (G) Representative data of GSIS assay in *FCHSD2*-KO cells. Data are mean ± SEM. *, *P* < 0.05; **, *P* < 0.01; ***, *P* < 0.005. *n* = 3. (H) Fold change of secreted insulin. Data are normalized to insulin secretion at basal level (0.5G). *, *P* < 0.05; **, *P* < 0.01; ***, *P* < 0.005. *n* = 4.

For STARD10 null EndoC-βH1 cells, we observed a significant reduction in glucose-stimulated insulin secretion in comparison with control (sham-transfected) cells [Fold change (15 mM glucose vs. 0.5 mM glucose): SHAM: 2.56 ± 0.14 vs. gRNA1: 2.12 ± 0.19, P = 0.007; vs. gRNA2: 1.70 ± 0.16, P = 0.007, *n* = 4] (Figure 6C and 6D), which is in agreement with our previous observations in mouse models (Carrat et al., 2017). Insulin secretion at low (0.5 mM) glucose was not affected by STARD10 loss. On the other hand, FCHSD2 null cells showed a relatively mild reduction of insulin secretion at both basal [Fold-change, 0.5 mM glucose vs. 0.5 mM glucose: SHAM: 1 ± 0 vs. gRNA1: 0.78 ± 0.053. P = 0.0265; vs. gRNA2: 0.66 ± 0.065 P = 0.013, *n* = 4] and high glucose conditions [Fold change, 15 mM glucose vs.15 mM glucose: SHAM: 1 ± 0 vs. gRNA1: 0.86 ± 0.016, P = 0.0043; gRNA2: 0.72 ± 0.01, P = 0.028, *n* = 4] (Figure 6G and 6H, Figure S6C and S6D). Taken together, these data demonstrate the involvement of STARD10 in insulin secretion in human β cells and unveil FCHSD2 as a previously unknown regulator of insulin exocytosis.

FCHSD2 has been shown to regulate F-actin polymerization suggesting it could be involved into insulin exocytosis (Cao et al., 2013). In order to identify any defect associated with FCHSD2 loss of function on exocytosis, we measured the membrane capacitance of FCHSD2 null cells using patch clamp electrophysiology. The membrane capacitance is proportional to the cell membrane surface area and is used as a readout of the exocytosis triggered by a train of ten depolarisations. On average, the maximal amplitude of stimulated exocytosis reached 94.3 ± 23.3 fF and 62.5 ± 17.0 fF for control (n=13) and FCHSD2 null (n=12) cells, respectively (Figure S6E). Normalised to the cell size, the cumulative increase in membrane surface area culminated to 9 ± 1.9 3 fF.pF^-1^ and 7.07± 2 fF.pF^-1^ in Control and FCHSD2 null cells respectively (student t-test at pulse 10, p=0.42) (Figure S6F).

We then focused on the kinetics of exocytosis by analysing the increment of membrane capacitance triggered by each pulse. In both lines, the increase of exocytosis was biphasic and mainly triggered by the two first pulses as previously described in EndoC-βH1 cell lines (Hastoy et al., 2018). The first pulse triggered a very similar amount of exocytosis in both groups (4.6 ± 0.8fF.pF^-1^ for control cells, and 4.7 ± 1.2 fF.pF^-1^ for FCHSD2 null cells) (Figure S6G). Therefore, the loss of FCHSD2 expression in EndoC-βH1 line does not affect significantly either the amplitude or the kinetics of exocytosis. In addition, no significant effects were observed on the size of the cells measured (Control: 9.9 ± 0.7pF, FCHSD2 null: 8.7 ± 0.7pF) (Figure S6H).

### Deletion of the variant region alters 3D chromatin structure and downregulates the expression of *STARD10* and *FCHSD2* genes

Finally, having examined the 3D structure and downstream genes of the enhancer cluster at the *STARD10* locus, we attempted to determine whether the risk-bearing variant region (VR) might affect enhancer cluster function. As shown in Figure 1A, the variant region is located between two active enhancers, R7 and R8 and, most importantly, is associated with CTCF binding regions R13 and R1 through chromatin looping (Figure 2C). These observations suggest that the VR may be involved in the formation of local chromatin structure and thus in controlling the activity of the enhancer cluster. To test this possibility, we performed 3C-qPCR analysis in sham control and dVR cells. In order to increase deletion efficiency of CRISPR-Cas9 editing in these experiments, we doubled the concentration of lentivirus (MOI = 20) and achieved ∼ 80 % (79.84 ± 0.59) deletion (Figure S7A-S7B). As shown in Figure 7A, VR deletion caused a significant change in 3D structure, notably a reduction in the physical interaction between the R13 and R1 regions (cross linking frequency: SHAM: 20.16 ± 1.144 vs. dVR: 13.3 ± 0.8525. P = 0.0064, *n* = 3). Further analysis of gene expression by Taqman qRT-PCR in dVR cells revealed that both the *STARD10* and *FCHSD2* genes were moderately down-regulated when compared with sham control cells (fold change: SHAM: 1.0 ± 0 vs. dVR: *STARD10*: 0.765 ± 0.03, P = 0.0159; *FCHSD2*: 0.859 ± 0.03, P = 0.011, *n* = 3). No significant change was observed in the expression of the *ARAP1* or *ATG16L2* genes (Figure 7B-7G). Taken together, these data demonstrate that the VR is an important region regulating chromatin 3D structure and transcriptional activity of the enhancer cluster.

**Figure 7.**
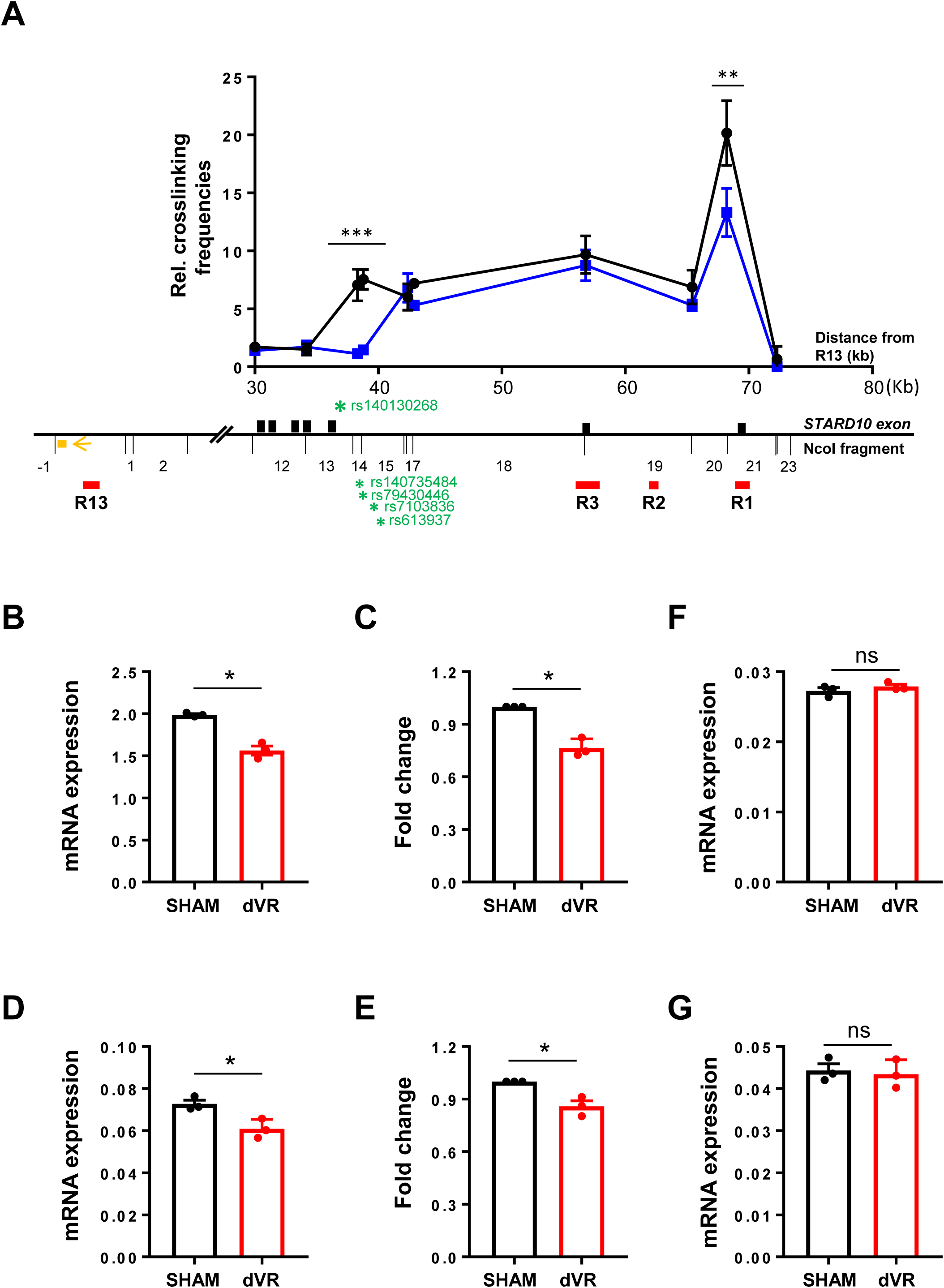
Effect of variant region deletion on 3D structure and gene expression. (A) Combined 3C-qPCR data of chromatin interactions in dVR cells. Black: control; Blue: dVR. The numbering of *NcoI* DNA fragments is given relative to the viewpoint. Viewing point: R13 region; Black bars: *STARD10* exon; red bars: regulatory region; orange box: qPCR probe; orange arrow: qPCR constant primer; green stars: credible set genetic variants. Data were normalized to a *CXCL12* loading control. *n* = 3. (B) Taqman qRT-PCR analysis of gene expression in control and dVR cells. *, *P* < 0.05; **, *P* < 0.01; ***, *P* < 0.005. *n* = 3.

## DISCUSSION

Our earlier study (Carrat et al., 2017) provided evidence, based on a combination of approaches (including human genetics and β-cell-selective gene knockout in the mouse), that *STARD10* is a critical mediator of the effects of the T2D-associated variants at this locus. However, our previous report did not explore in detail how the identified credible set may influence the expression of local genes, nor did it exclude the possibility that other genes might also be involved in the actions of risk variants.

The goals of the present study were, therefore, to obtain a more detailed molecular picture of the local chromatin structure at the *STARD10* T2D locus, and to use this to identify and study genes likely to interact with previously identified T2D risk variants in human β cells. In this way, we report a likely spatial organization, defined by CTCF-stabilized looping, which allows an enhancer cluster to regulate not only the *STARD10* gene, but also the distal *FCHSD2* gene, which is contained in the same 3D chromatin compartment as *STARD10* and the T2D risk variants. In contrast, and consistent with our earlier studies (Carrat et al., 2017), we found no evidence for a role for *ARAP1* in mediating the effects of the variants on β cell function.

We also report here that variants of the risk haplotype are eQTLs for both *STARD10* and *FCHSD2* in human islets (note that data were not available for the indel with highest posterior probability). Importantly, these results were only obtained in laser capture microdissection (LCM) donors from partially pancreatectomized patients from the “IMIDIA” data set (Carrat et al., 2017). In contrast, interrogation of OD samples in the same dataset fail to reveal a nominal association between any of the T2D risk SNPs and *FCHSD2* expression. Moreover, interrogation of other islet eQTL datasets reported earlier, which are also derived from OD samples (Mahajan et al., 2018)(Miguel-Escalada et al, 2019), provided no evidence of association with islet *FCHSD2* levels.

We note that the use of LCM to extract mRNA is unlike enzymatic digestion methods commonly used with OD samples. Thus, surgical specimens are subject to immediate cryofixation, limiting RNA degradation and hence transcriptomic changes which can occur in OD samples where islets are isolated from the rest of the pancreas prior to mRNA extraction. Moreover, LCM methods provide a purer and more β cell-enriched cell population (Solimena et al., 2018). Correspondingly, OD and LCM samples from the same individual cluster separately (Solimena et al., 2018). Thus, the detection of the above eQTL for *FCHSD2* in the IMIDIA cohort would appear most likely to be a function of the availability in this cohort of mRNA extracted using LCM.

We used a CRISPR-Cas9 approach to delete the whole of the 4.1 Kb variant region likely to host the credible set of T2D variants with significant posterior probability at this locus (Carrat et al., 2017). This region was also identified as a credible set in earlier reports (Figure S5C) (Gaulton et al., 2015 and Bonàs-Guarch et al., 2018). We note, however, that it did not feature in a recent report (Mahajan et al., 2018). Unlike our previous analysis, the new report from Mahajan *et al*. does not include indels [a Haplotype Reference Consortium (HRC) panel was used which only included SNPs]. Thus, the variant in the VR region with the highest PPA (indel rs140130268), as described in our earlier report (Carrat et al., 2017) in which a Metabochip scaffold was imputed up to 1000G reference panel including indels, was not included. We note that whereas indel rs140130268 showed a PPA of 45 % (Carrat et al., 2017), in Mahajan *et al*, maximum PPA is assigned to rs7109575. These two SNPs are in very high LD (EUR R2 = 0.95). In any case, the discordance between earlier studies emphasizes the need for interventional studies, as presented in this report, in which the impact of deleting the implicated regions is explored directly, at a functional level, in disease-relevant cells.

Whilst the ideal design of such studies would involve using homologous recombination to convert risk into protective alleles (or *vice versa*) one by one, currently available cellular systems largely preclude this. EndoC-βH1 β cells grow slowly and do not tolerate cloning through a single cell stage, whilst human embryonic stem-cell (ES)-derived β cells (Pagliuca et al., 2014; Rezania et al., 2014 and Nair et al., 2019) do not reliably provide a robust platform for functional studies, notably of insulin secretion.

Enhancer hubs contain genes that are key for the regulation of β cell function, and that the activity of elements (genes and enhancers) of the same hub tends to be correlated (Miguel-Escalada et al., 2019). Although the approach adopted here to delete the whole 4.1 kb region in EndoC-βH1 β cells provides no information on the role of the *individual* variants, it does, nonetheless, allow us to suggest that this region is important for the overall regulation of the enhancer hub, and therefore in the control of local gene expression and β cell function.

### Enhancer hub and chromatin structure at the *STARD10* locus

The spatial organization of chromatin can play an important role in gene regulation (Vernimmen and Bickmore, 2015). We found that the T2D risk variants at the *STARD10* locus, reside in an islet enhancer hub which contains the genes *ARAP1, STARD10, ATG16L2* and *FCHSD2*. Of note, expression of *ARAP1* in human islets is chiefly driven by two promoters that reside outside the enhancer hub (Figure 5A and Figure S5C), and is thus not likely to be co-regulated with *STARD10* and other hub genes, in line with evidence from eQTL studies (this report and (Carrat et al., 2017)).

Using CRISPR-Cas9 and ChIP analyses, we further determined that the two CTCF-binding sites located at either end of the cluster contribute to the functional integrity of the hub. Together with a strong enhancer (R2) and binding of β cell-specific transcriptional factors (Figure 1A), this creates a spatially organized transcriptional complex within which variants that confer differential regulatory potential influence the expression of relevant genes.

### *STARD10* and *FCHSD2* mediate altered β cell function and disease risk

Our previous studies (Carrat et al., 2017) provided functional evidence, through gene deletion, of an essential role for *STARD10* in mediating glucose-induced insulin secretion, and proinsulin processing, in the mouse. Here, using CRISPR-Cas9-mediated inactivation, we demonstrate that STARD10 is equally important in human β cells. Although the molecular roles for STARD10 in the β cell remain to be elucidated, our preliminary findings (Carrat et al, submitted) indicate a role in the control of secretory granule biogenesis, presumably reflecting a requirement for STARD10-mediated intracellular lipid transport between relevant donor and acceptor membranes. Of note, we did not, in the present studies, explore the impact of *STARD10* deletion on proinsulin processing. When assessed through plasma proinsulin:insulin ratios this is decreased in risk variant carriers (Strawbridge et al., 2011) and *STARD10* KO mice (Carrat et al., 2017). However, the difference is not readily detected *in vitro* comparing islets from wild-type and null mice.

Analysis of human islet pcHiC (Miguel-Escalada et al., 2019), as well as measurements of changes in gene expression after deletion of R2 region or the core of the variant region, now provide evidence that *FCHSD2* may also play a role in mediating disease risk. Interestingly, deletion of the variant region led to a lowering of stimulated insulin secretion, but no change in basal insulin secretion (Figure 1F and 1G). Whereas knockout of *FCSHD2* lowered both basal and stimulated insulin release (Figure 6H and Figure S6C), loss of *STARD10* had no effect on insulin secretion at low glucose, but impaired insulin secretion at high glucose (Figure 6C and 6D). It might therefore be speculated that the variant region exerts effects on insulin secretion via a combined action on both genes. Future studies will be needed to explore this possibility in more detail.

By what mechanisms may FCHSD2 affect insulin secretion? FCHSD2 has been shown to regulate F-actin polymerization (Cao et al., 2013). More recently, it has also been found to act as a positive regulator of clathrin-mediated endocytosis (Almeida-Souza et al., 2018). FCHSD2 contains four distinct domains: an N-terminal F-BAR domain containing an atypical additional coiled coli (CC) at its C terminus, two SH3 (src homology 3) domains and a C-terminal proline rich region (PRR). FCHSD2 is recruited to clathrin-coated pits by intersection through its second HS3 domain (SH3-2), while its first SH3 domain (SH3-1) binds to N-WASP to initiate F-actin polymerization (Almeida-Souza et al., 2018 and Xiao et al., 2018). A role for FCHSD2 in insulin exocytosis has not previously been considered. We note that exocytosis requires F-actin-mediated cytoskeletal remodeling (Kalwat and Thurmond, 2013 and Jewell et al., 2008) in which N-WASP and ARP2/3 complex are important protein complex in the formation of focal adhesion. It is therefore conceivable that, through this mechanism, FCHSD2 plays an active role in the regulation of glucose-mediated insulin exocytosis. However, although a slight tendency was observed towards lowered exocytotic capacity in FCHSD2 null cells, our initial results have not, so far, provided firm evidence in support of such a role (Figure S6E-S6H). It is also conceivable that FCHSD2 plays a role in controlling the localization at the plasma membrane, or recycling, of receptors, such as that for glucagon-like peptide-1 (GLP1), which are important in the physiological responses to insulin secretion and food intake (Jones et al., 2018).

### Possible impact of T2D variants on chromatin landscape at the *STARD10* locus

Although not explored directly in the present study, we note that the VR region is located between two enhancers (R8 and R7) which are occupied by β-cell specific TFs such as NKX2.2, PDX1 and MAFB (Figure 1A). The VR itself also displayed strong PDX1 binding (Figure 1A), consistent with a β cell-specific role within enhancer cluster. We demonstrate here that the VR interacts with CTCF binding regions and its deletion causes a significant change in 3D structure, affecting the expression of downstream genes *STARD10* and *FCHSD2* (Fig. 7).

As far as we are aware, this is therefore the first study to demonstrate directly a change in chromatin structure as a result of deleting a genomic region hosting a variant associated with altered T2D risk. Future studies, in more tractable systems (e.g. CRISPR-edited human embryonic stem cell-derived β-like cells) are likely to be required to determine the effect of more targeted (e.g. single variant or haplotype) changes in DNA sequence.

In summary, the present report extends our previous studies of the *STARD10* locus, revealing important aspects of the local chromatin structure and identifying a functionally relevant new gene at this locus. Our studies also demonstrate that the T2D-associated variant region may influence local chromatin structure to alter the expression of a nearby gene embedded within an enhancer cluster (*STARD10*), as well as that of a more remotely located gene (*FCHSD2*) through a long-range chromatin loop.

## ACKNOWLEDGMENTS

G.A.R. was supported by Wellcome Trust Senior Investigator (WT098424AIA) and Investigator (212625/Z/18/Z) Awards, MRC Programme grants (MR/R022259/1, MR/J0003042/1, MR/L020149/1) and Experimental Challenge Grant (DIVA, MR/L02036X/1), MRC (MR/N00275X/1), Diabetes UK (BDA/11/0004210, BDA/15/0005275, BDA 16/0005485) and Imperial Confidence in Concept (ICiC) grants, and a Royal Society Wolfson Research Merit Award. BH thanks Diabetes UK for an R.D. Lawrence Fellowship (BDA:19/0005965). This project has received funding from the European Union’s Horizon 2020 research and innovation programme via the Innovative Medicines Initiative 2 Joint Undertaking under grant agreement No 115881 (RHAPSODY) to G.A.R., P.M., P.F., and M.S. This Joint Undertaking receives support from the European Union’s Horizon 2020 research and innovation programme and EFPIA. Finally, we thank Jorge Ferrer for inputs and helpful discussion.

## AUTHOR CONTRIBUTIONS

M.H. designed and performed all molecular and cellular experiments (EMSA, promoter-luciferase assay, CRISPR-Cas9 editing, ChIP-qPCR and insulin secretion analyses). I.C. provided and analyzed epigenomic and promoter Hi-C data. G.C. performed Western blot analysis. S.J. generated the CRISPR constructs for *STARD10* under M.H. supervision. A.K., M.C., P.F., A.S., M. S., M.I., P.M., L.C. provided human islet samples and performed eQTL analysis. P.G. provided the β cell-specific lentiviral vector for CRISPR-Cas9 targeting. L.A. and H.McM. provided human FCHSD2 antibody. G.R. conceived the project with M.H. and supervised all studies. M.H., I.C., and G.R. wrote the manuscript with input from all authors.

## DECLARATION OF INTERESTS

G.R. has received grant funding from Les Laboratoires Servier and is a consultant for Sun Pharmaceuticals.

## STAR Methods

### METHOD DETAILS

#### Cell Culture

The human-derived β cell line EndoC-βH1 was grown on ECM (1% v/v) and Fibronectin (2 μg/ml)-coated plates or petri dishes in serum-free DMEM containing low glucose (1 g/L), 2 % (w/v) albumin from bovine serum fraction V, 50 μM β-Mercaptoethanol, 10 mM nicotinamide, 5.5 μg/mL human transferrin, 6.7 ng/mL sodium selenite, penicillin (100 units/mL), and streptomycin (100 μg/mL) (Ravassard et al., 2011). HEK293T cell was cultured in DMEM high glucose medium supplemented with 10 % fetal bovine serum, 6 mM L-glutamine, penicillin (100 μg/mL) and streptomycin (100 μg/mL).

#### Electrophoretic Mobility Shift Assay (EMSA)

Electrophoretic mobility shift assay (EMSA) was carried out as previously described (Stewart et al., 1997). In brief, complementary oligonucleotides were designed to contain either risk or protective variants (Table S3). EndoC-βH1 nuclear extract was prepared using NE-PER nuclear and cytoplasmic extraction kit according to manufacturer’s instruction (Thermo Scientific). Oligonucleotides bearing either risk or protective variants were synthesized (Sigma) and end-labelled with γ-^32^P-ATP using T4 polynucleotide kinase (New England BioLabs). ^32^P-labelled oligoes were incubated for 20 min at room temperature with 5 μg of nuclear extract in binding reactions consisted of 1 x binding buffer (20 mM Hepes pH 7.9, 90 mM KCl, 5 mM MgCl_2_, and 0.05 % Nonidet P-40) and 1 μg poly(dI-dC). Samples were electrophoresed on a 5 % acrylamide gel in 0.5 x TBE buffer (90 mM Tris, 64.6 mM Boric acid and 2.5 mM EDTA, pH 8.3). The acrylamide gel was then vacuum-dried and autoradiographed.

#### Chromatin Conformation Capture (3C and 4C)

3C was performed as described (Hagège et al., 2007). In brief, a suspension of 1 × 10^7^ EndoC-βH1 cells was cross-linked with 4 % (v/v) formaldehyde at room temperature for 10 min. The cross-linked DNA was digested overnight with restriction enzyme *Nco*I and then ligated with T4 DNA ligase at 16° C overnight. The ligated 3C DNA was purified by extraction with phenol/chloroform and precipitation with ethanol. The ligation products were quantitated by Taqman™ qPCR and normalized to the human *CXCL12* gene. The standard curve for each primer pair was generated using *Nco*I-digested and ligated BAC DNA (RP11-101P7) encompassing the human *ARAP1, STARD10*, and *ATG16L2* loci. The Taqman™ probes and primers used for the 3C experiments presented in this study are listed in Table S8. Taqman™ qPCR were carried out on a 7500 Fast Real-Time PCR System (Applied Biosystems). PCR reactions were set as follows: 95°C for 10 min., then with 45 cycles at 95°C 30 s and 58°C 45 s. Crosslinking frequencies were plotted as percentage of that of the human *CXCL12* gene. *Bam*HI-digested and ligated 3C sample was used as a negative control.

For 4C, 1 × 10^7^ EndoC-βH1 cells were fixed with 4 % paraformaldehyde, digested with restriction enzyme *Pst*I, ligated with T4 DNA ligase and then digested with second restriction enzyme *Msp*I. After the second round of DNA ligation with T4 DNA ligase, DNAs were purified with phenol/chloroform extraction and ethanol precipitation. PCR reactions were carried out to amplify ligation products using nested PCR primer sets (Table S7). PCR products were then digested with restriction enzymes *Xho*I and *Not*I and sub-cloned into pBluescript II KS+ (pBSKS) (Stratagene/Agilent) for Sanger sequencing analysis.

#### Chromatin Immunoprecipitation

Immunoprecipitation was carried out according to a standard protocol (Boj et al., 2001). In brief, 1 × 10^6^ EndoC-βH1 cells were fixed with 1 % (v/v) formaldehyde for 10 min. and quenched with 1.25 mM glycine. Cells were then scraped and resuspended in lysis buffer (2 % Triton-100, 1 % SDS, 100 mM NaCl, 10 mM Tris-HCl, 1 mM EDTA). After 20 stokes of homogenization with a disposable pestle, cells were sonicated for 10 min. using Covaris™ S220 to breakdown genomic DNA to 200-500 bp fragments. DNA/protein complexes were then precipitated with anti-CTCF antibody or rabbit IgG (EMD Millipore) conjugated with protein A and G beads. DNAs were purified through Phenol/Chloroform extraction and Ethanol precipitation.

#### PCR and qPCR

Fusion high fidelity Taq polymerase (Thermo Fisher Scientific) was used in all routine PCR reactions to avoid PCR errors. A typical PCR reaction was set as follow: 98°C for 30 s, then with 35 cycles at 98°C 10 s, 60°C 10 s and 72°C 15 s. The primer sets for genomic DNA amplification are listed in Table S5.

Total RNA from EndoC-βH1 cells was obtained using TRIzol reagent (Invitrogen). Total RNAs (2 μg) were then reverse-transcribed into first strand cDNAs using High-Capacity cDNA Reverse Transcription Kit (Thermo Fisher Scientific) according to the manufacturer’s instructions. Real-time PCR was performed on a 7500 Fast Real-Time PCR System using the Fast SYBR™ Green master mix or Fast Taqman™ master mix. The SYBR™ Green PCR primer sets for variant region and CTCF binding site (CBS) are listed in Table S6 and Table S9, respectively.

#### Molecular Cloning

Active enhancer regions, identified by integration of previously published human islet ATAC-seq and H3K27ac ChIP-seq datasets (Miguel-Escalada et al., 2019), were PCR-amplified from BAC DNA (RP11-101P7) with primer sets (Table S10) designed by Primer3-based software and cloned into pGL3-promoter vector between NheI and XhoI restriction enzyme sites. Plasmid DNA was extracted using mini-prep plasmid extraction kit and/or Maxi-prep plasmid extraction kit (Qiagen). Correct cloning was confirmed by Sanger sequencing.

#### Transfection and Luciferase Assay

EndoC-βH1 cells were seeded at a density of 50,000 per well in 48-well plates. After 48 h, 0.4 μg of luciferase constructs containing putative regulatory sequences were co-transfected with 1 ng of pRL-Renilla construct as internal control into EndoC-βH1 cells, using Lipofectamine 2000, according to manufacturer’s instruction. pGL3-promoter vector was served as a negative control. 48 h later, transfected cells were washed once with PBS and lysed directly in passive cell lysis buffer (Promega). Cells were incubated on a rotating platform at room temperature for 10 min. to ensure complete lysis of cells, and then spun at 10,000 rpm for 10 min to remove cell debris. Supernatant was transferred into a fresh tube and used to measure luciferase activity with Dual-Luciferase Reporter Assay kit (Promega) on a Lumat LB9507 luminometer (Berthold Technologies). Firefly luciferase measurements were normalized to *Renilla* luciferase.

#### CRISPR-Cas9-mediated Genome Editing

gRNA sequences were designed using the software provided by Broad Institute (https://portals.broadinstitute.org/gpp/public/analysis-tools/sgrna-design) (Doench et al., 2016). To generate mutations or deletions in EndoC-βH1 cells, lentiviral constructs carrying both gRNA and humanized *S. pyogenes* Cas9 (*hsp*Cas9) were transfected into HEK293T cells together with packaging plasmids PMD2.G and psPAX2 using CaCl_2_ transfection protocol (Graham and van der Eb, 1973). The lentiviral vector containing RIP-Cas9 gene cassette but without gRNA was served as a SHAM control. Next day, cells were treated with sodium butyrate (10 mM) for 8 hours before changing to fresh medium. The medium was collected twice in the next two days and subjected to ultracentrifugation (Optima XPN-100 Ultracentrifuge, Beckman Coulter) at 26,000 rpm for 2 hours at 4°C. The lentiviruses were collected from the bottom of the tube and titrated. Same amount of viruses were used to transduce to EndoC-βH1 cells (MOI = 10). Puromycin (4 μg/ml) was added 72 h after infection to select lentivirus-infected cells. For deletion of genomic regions, two plasmids carrying two different gRNAs flanking target regions were co-transfected into HEK293T cells with packaging plasmids. All sequences of gRNAs and primers used for genotyping of genome editing experiments in this study are listed in Table S4 and Table S5, respectively.

To measure deletion efficiency after CRISPR/Cas9 mediated genome editing, Cyber Green qPCR was deployed to detect wildtype allele using primers 1 and 2 from genomic DNA extracted from control and VR-deleted or R2-deleted cells. *CXCL12* was served as an internal DNA copy number control (key resources table). The deletion efficiency was calculated as: [1-2^ΔΔCt(del-CXCL12)^/2^ΔΔCt(WT-CXCL12)^] x 100%. In addition, the relative values of DNA deletion or inversion in VR or R2 deleted cells was also measured using primer set 1+4 or 1+3 respectively. The primers for VR-del and R2-del were listed in Supplementary Table S6 and S11, respectively.

#### Electrophysiology

Membrane capacitance measurements were performed at 37°C in standard whole cell configuration. The recording were made using an EPC-10 amplifier and Pulse software as previously described. In brief, the extracellular medium was composed of (mmol/L) 118 NaCl, 3.6 KCl, 0.5 MgSO_4_, 0.5 NaH_2_PO_4_, 5 NaHC0_3_, 1.5 CaCl_2_, 5 HEPES, and 20 tetraethylammonium (TEA) (pH 7.4 with NaOH). The intracellular medium contained (mmol/L) 129 CsOH, 125 Glutamic acid, 20 CsCl, 15 NaCl, 1 MgCl_2_, 0.05 EGTA, 3 ATP, 0.1 cAMP, 5 HEPES (pH7.2 with CsOH). The cells were subjected to ten 500ms depolarisations from - 70mV to 0mV at 1Hz. Data were analysed using R software and plotted in Prism software. These measurements were performed over three independent passages of EndoC-βH1 CRISPR Control (n=13 cells) and of FCHSD2 CRISPR KO (n=12 cells) lines. Data are expressed as mean±sem with the detailed data point overlapped.

#### Insulin Secretion

EndoC-βH1 cells were seeded onto ECM/Fibronectin-coated 48-well plates at 2.5 × 10^5^ cells per well. Four days after seeding, cells were incubated overnight in a glucose starving medium (glucose-free DMEM supplemented with 2 % Albumin from bovine serum fraction V, 50 μl 2-mercaptoethanol, 10 mM nicotinamide, 5.5 μg/ml transferrin, 6.7 ng/ml sodium selenite, 100 units/ml, penicillin, 100 μg/ml streptomycin and 2.8 mM glucose).

The next morning cells were incubated for 1 h in Krebs-Ringer solution [0.2 % BSA, 25 % solution 1 (460 mM NaCl), 25 % solution II (96 mM NaHCO_3_, 20 mM KCl and 4 mM MgCl_2_), 25 % solution III (4 mM CaCl_2_), 10 mM Hepes] supplemented with 0.5 mM glucose. EndoC-βH1 cells were then incubated in the presence of low (0.5 mM) or high glucose (15 mM) or other stimuli [0.5 mM IBMX or 20 nM Forskolin or 20 mM KCl]. After incubation for 1 h, the supernatant was collected, placed onto ice and centrifuged for 5 min. at 3,000 rpm at 4 °C. The supernatant was then transferred into a fresh tube. Cells were lysed in 50 μL of cell lysis solution (TETG: 20 mM Tris pH 8.0, 1 % Triton X-100, 10 % glycerol, 137 mM NaCl, 2 mM EGTA). The lysate was then removed to a fresh tube and centrifuged at 3,000 rpm for 5 min at 4°C. Insulin content was measured using an insulin ultra-sensitive assay kit. Secreted insulin was normalized as percentage of total insulin content. Fold increase in glucose- or other stimuli-stimulated insulin secretion is expressed as a ratio in comparison with secretion at basal level (0.5 mM glucose).

#### Human Islet Regulome and Interactome Analysis

Human islet regulome maps, including accessible chromatin regions (ATAC-seq) and ChIP-seq for H3K27ac, CTCF and different islet-enriched TFs (NKX2.2, FOXA2, PDX1 and MAFB) were obtained from previously published datasets (Pasquali et al., 2014 and Miguel-Escalada et al., 2019) and visualized using the UCSC Genome Browser (http://genome.ucsc.edu/) with GRCh37/hg19 assembly. Represented data corresponds to consolidated tracks released by Miguel-Escalada et al. Details on number of samples per track are available (Miguel-Escalada et al., 2019). Previously published human islet RNA-seq (Morán et al., 2012), chromatin interaction maps (Miguel-Escalada et al., 2019a) were visualized on the WashU Epigenome browser using this session link: http://epigenomegateway.wustl.edu/browser/?genome=hg19&session=62hGf7nfcS&statusId=140947077. For pcHi-C interactions, only interactions with a CHiCAGO score > 5 were taken as high-confidence interactions, as previously described (Miguel-Escalada et al., 2019). For visualization of purposes only, pcHi-C interactions mediated by the CTCF sites R1 and R13 were colored in orange and all other high-confidence interactions in grey (Figure 5A).

#### Transcription Factor Binding Motif Analysis

TF binding profile on genetic variants was carried out using JASPAR CORE program (http://jaspar.genereg.net) (Khan et al., 2018). The threshold of relative profile score was set up at 80%. The scores of potential transcription factors were compared between risk and protective variants and listed in Table S2.

#### Expression Quantitative Trait Loci (eQTL) Analysis

Pancreatic tissues and blood samples were collected from 103 patients that have undergone partial pancreatectomy from the IMIDIA consortium (Khamis et al., 2019 and Solimena et al., 2018), with appropriate permissions from donors and/or families. Briefly, expression data was acquired from islets isolated by laser capture microdissection from surgical specimens using Human Genome U133 Plus2.0 Array (Affymetrix). DNA was genotyped using the 2.5 M Omniarray beadchip (Illumina) and imputed with 1,000 Genomes reference panel (phase 3), resulting in 7.5 M SNPs. Standard quality control assessment was carried out on the genotyping data using PLINK (Purcell et al., 2007). Expression and genotype analysis was combined to generate eQTLs, performed with FastQTL (Ongen et al., 2016) with gender and age as covariates. A cis-window of 500 kb was used, *i.e.*, the maximum distance at which a gene-SNP is considered local.

#### Statistical Analysis

Data are expressed as means ± SEM. Significance was tested by Student’s two-tailed t test, Mann-Whitney test for non-parametric data, and one- or two-way ANOVA with SIDAK multiple comparison test, as appropriate, using Graphpad Prism 7.0 software. P<0.05 was considered significant.

