## Supplemental Methods for "Chromatin 3D interaction analysis of the *STARD10* locus unveils *FCHSD2* as a new regulator of insulin secretion"

**STAR Methods**

Key Resources Table

| REAGENT or RESOURCE | SOURCE | IDENTIFIER |
|---|---|---|
| **Antibodies** | | |
| CTCF | EMD Millipore | Cat# 07-729; RRID: AB_441965 |
| Rabbit IgG, plasma | EMD Millipore | Cat# 401590; RRID: N/A |
| STARD10 | Santa Cruz Biotechnology | Cat# sc-54336; RRID: AB_2197780 |
| FCHSD2 | (Almeida-Souza et al., 2018) | N/A |
| **Bacteria and Virus Strain** | | |
| Stbl3 competent cells | Thermo Fisher Scientific | Cat# C737303 |
| 10-beta competent cells | New England Biolabs | Cat# C30191 |
| **Biological samples** | | |
| Human islet samples | (Carrat et al., 2017) | N/A |
| **Chemicals, Peptides, and Recombinant Proteins** | | |
| 3-Isobutyl-1-methylxanthine (IBMX) | Sigma-Aldrich | Cat# I5879 |
| Foskolin | Sigma-Aldrich | Cat# F3917 |
| Sodium Selenite | Sigma-Aldrich | Cat# S1382 |
| Nicotinamide | Sigma-Aldrich | Cat# 481907 |
| Paraformaldehyde | Agar Scientific | Cat# AGR1026 |
| TRIzol | Invitrogen | Cat# 15596018 |
| NP40 | Sigma-Aldrich | Cat# 13021 |
| Proteinase inhibitor cocktail | Sigma-Aldrich | Cat# 04693132001 |
| Albumin from Bovine Serum Fraction V | Roche Diagnostics | Cat# 10775835001 |
| Human Transferrin | Sigma-Aldrich | Cat# T8158 |
| ECM | Sigma-Aldrich | Cat# E1270 |
| RNase A | Thermo Fisher Scientific | Cat# EN0531 |
| Proteinase K | Sigma-Aldrich | Cat# AM2546 |
| Dynabeads-Protein G | Thermo Fisher Scientific | Cat# 10003D |
| Dynabeads-Protein A | Thermo Fisher Scientific | Cat# 10008D |
| ATP, [γ-$^{32}$P] | Perkin Elmer | Cat# NEG002A |
| **Critical Commercial Assay** | | |
| Dual-Luciferase Assay Reporter Assay System | Promega | Cat# E1910 |
| Lipofectamine 2000 | Thermos Fisher Scientific | Cat# 11668025 |
| Insulin Ultra-sensitive Assay Kit | Cisbio Bioassays | Cat# 62IN2PEH |
| High-Capacity cDNA Reverse Transcription Kit | Thermos Fisher Scientific | Cat# 4368814 |
| NE-PER$^{TM}$ Nuclear and Cytoplasmic extraction reagents | Thermos Fisher Scientific | Cat# 78833 |
| Taqman$^{TM}$ Fast advanced master mix | Thermos Fisher Scientific | Cat# 4444553 |
| Fast SYBR$^{TM}$ Green Master Mix | Thermos Fisher Scientific | Cat# 4385612 |

| | | |
|---|---|---|
| Phusion High fidelity DNA polymerase | Thermos Fisher Scientific | Cat# F530 |
| T4 polynucleotide kinase | New England BioLabs | Cat# M0201S |
| **Experimental Model: cell line** | | |
| HEK293T | ATCC | Cat# CRL_3216; RRID: CVCL_0063 |
| EndoC-$\beta$H1 | Univercell-Biosolutions | Cat# N/A; RRID: CVCL_L909 |
| **Oligonucleotides** | | |
| EMSA assay | See Table S3 | |
| gRNAs | See Table S4 | |
| Genomic DNA amplification | See Table S5 | |
| SYBR$^{TM}$ green qPCR for VR region | See Table S6 | |
| 4C assay in VR region | See Table S7 | |
| 3C assay in VR region | See Table S8 | |
| SYBR$^{TM}$ green qPCR for CBSs | See Table S9 | |
| 3C assay in *STARD10* locus | See Table S8 | |
| Luciferase assay | See Table S10 | |
| **Recombinant DNA** | | |
| pMD2.G | Didier Trono (https://tronolab.epfl.ch) | Cat# 12259; RRID: Addgene_12259 |
| pSPAX2 | Didier Trono (https://tronolab.epfl.ch) | Cat# 12260; RRID: Addgene_12260 |
| BAC DNA | bacpacresources.org | RP11-101P7 |
| pLenti-CRISPR-RIP-Cas9 | Cardenas-Dias et al., Cell Stem Cell, accepted in principle | N/A |
| pGL3-promoter | Promega | E1761 |
| pRL-Renilla | Promega | E2231 |
| pBlueScript II KS+ | Stratagene/Agilent | 212207 |
| **Taqman$^{TM}$ qPCR primer/probe** | | |
| *CXCL12* | Thermos Fisher Scientific | Cat# 4400291; Assay ID: Hs02117611_cn |
| *STARD10* | Thermos Fisher Scientific | Cat# 4331182; Assay ID: Hs00246405_ml |
| *ARAP1* | Thermos Fisher Scientific | Cat# 4331182; Assay ID: Hs00362929_ml |
| *FCHSD2* | Thermos Fisher Scientific | Cat# 4331182; Assay ID: HS00207952_ml |
| *ATG16L2* | Thermos Fisher Scientific | Cat# 4331182; Assay ID: Hs01057324_ml |
| *ACTB* | Thermos Fisher Scientific | Cat# 4331182; assay ID: Hs01060665_gl |
| **Software** | | |
| GraphPad Prism 7 | GraphPad Software | RRID: SCR_002798 |
| Illustrator | Adobe | RRID:SCR_010279 |
| Photoshop | Adobe | RRID: SCR_014199 |

### SUPPLEMENTARY INFORMATION

#### Supplementary Figure Legends

**Figure S1. Credible set T2D variants at *STARD10* locus.**
(A) Potential transcription factor (TF) bindings at risk or protective variants. The binding affinity of TFs was analyzed and scored using JASPAR program (http://jaspar.genereg.net/). Red: risk variants; blue: protective variant in blue.
(B) Sanger sequencing of PCR product amplified from wild-type and dVR cell. Orange and blue bars: 5' and 3' end of DNA sequences flanking VR region, respectively.
(C) Diagram of SYBR$^{TM}$ Green qPCR analysis at VR region. Primers 1 and 2 were designed to amplify wildtype genomic DNA; primers 1 and 3 detect inversion of DNA after editing and primers 1 and 4 amplify DNA fragment after deletion. CXCL12 gene was served as an internal DNA copy number control.
(D) Representative data of SYBR$^{TM}$ Green qPCR analysis on wildtype and dVR genomic DNAs. Note that the rate of DNA inversion detected by primer 1+3 was about 10% of remaining wildtype genomic DNA (primer 1+2) in dVR cells, roughly count about 5% of efficiency rate when compared with wildtype control.
(E) Deletion efficiency. The deletion efficiency rate (%) was determined by: 1. calculating the remaining wildtype allele (primer 1+2) in dVR genomic DNA and compared with wildtype allele in SHAM control and 2. Taking away the rate of inversion (primer 1+3) in dVR cells. Data are mean ± SEM. *, $P < 0.05$; **, $P < 0.01$; ***, $P < 0.005$. N = 3.

**Figure S3. CTCF-binding site (CBS) at *STARD10* locus.**
(A) DNA sequencing diagrams of ChIP-qPCR identified CBSs at R13 and R1 regions. CBS1 and 2 are within R13, CBS5 is 2.7 kb downstream of R13, and CBS7 and 8 are within R1 region.
(B) Comparison of CTCF binding affinity before and after CRISPR-Cas9 editing at CBSs. CTCF binding affinity was normalized to IgG control in wild-type and CBS mutate cells. The numbers above each bar represent the fold change before and after genome editing. The data are pulled from two independent experiments.
(C) Representative data of Insulin secretion stimulated by multiple stimuli. Cells were treated with 0.5 mM glucose and then stimulated with either IBMX (0.5 mM) or Forskolin (20 nM) or KCl (20 mM). Data are mean ± SEM. *, $P < 0.05$; **, $P < 0.01$; ***, $P < 0.005$. N = 3.
(D) Fold change of secreted insulin. Data are normalized to insulin secretion at basal level (0.5G). *, $P < 0.05$; **, $P < 0.01$; ***, $P < 0.005$. N = 3.
(E) Representative data of Taqman$^{TM}$ qRT-PCR analysis in CBS mutate cells. G. *ARAP1*; H. *STARD10*; I. *ATG16L2* and J. *FCHSD2*. *, $P < 0.05$; **, $P < 0.01$; ***, $P < 0.005$. N = 3.

**Figure S4. Location and DNA sequence of R2 enhancer.**
(A) R2 enhancer is located between two promoters of *STARD10* gene (R3 and R1) with multiple binding sites for islet-associated transcription factors.
(B) Diagram of primers designed for SYBR$^{TM}$ Green PCR analysis at R2 region. Primers 1 and 2 were designed to amplify wildtype genomic DNA; primers 1 and 3 were to detect DNA inversion after genome editing and primers 1 and 4 were to amplify DNA fragment after deletion. CXCL12 gene was served as an internal DNA copy number control.
(C) Representative data of SYBR$^{TM}$ Green qPCR analysis in SHAM and dR2 cells.

(D) Deletion efficiency. The deletion efficiency rate (%) was determined by: 1. calculating the remaining wildtype allele (primer 1+2) in dR2 genomic DNA and compared with wildtype allele in wildtype control and 2. Taking away the rate of inversion (primer 1+3) in dR2 cells. Data are mean ± SEM. *, $P < 0.05$; **, $P < 0.01$; ***, $P < 0.005$. N = 2.

**Figure S5. Taqman$^{TM}$ qRT-PCR analysis of Gene expression in dR2 EndoC-βH1 cells**
(A) Representative data of *ARAP1* gene expression in dR2 cells. *, $P < 0.05$; **, $P < 0.01$; ***, $P < 0.005$. N = 3.
(B) Representative data of *ATG16L2* gene expression in dR2 cells. *, $P < 0.05$; **, $P < 0.01$; ***, $P < 0.005$. N = 3.
(C) Genome browser view of the *ARAP1/STARD10* locus displaying the three putative promotors for *ARAP1* (P1-3). Human islet RNA-seq track shows that exons belonging to *ARAP1* isoforms expressed from P1 and P2 are very lowly expressed in comparison to the isoform driven by the P3 promoter. We have modified the scale range to highlight this better. Please note that only P1 belongs to the islet enhancer hub (hub elements track). High PP SNPs track shows all T2D credible set SNPs with PP > 0.05 (Carrat et al., 2017).

**Figure S6. Knockout effects of *STARD10* or *FCHSD2* in EndoC-βH1 cells.**
(A) Sanger sequencing of PCR products amplified from wild-type and *STARD10*-KO cells. Red box: PAM sequence; red arrow: Cas9 cutting site.
(B) Sanger sequencings of PCR products amplified from wild-type and *FCHSD2*-KO cells. Red box: PAM sequence; red arrow: Cas9 cutting site.
(C) GSIS assay of *FCHSD2*-KO cells. Fold change at high glucose level (15G:15G). Data are mean ± SEM. *, $P < 0.05$; **, $P < 0.01$; ***, $P < 0.005$. N = 4.
(D) GSIS assay of *FCHSD2*-KO cells. Fold change at 15G vs. 0.5G. Data are mean ± SEM. *, $P < 0.05$; **, $P < 0.01$; ***, $P < 0.005$. N = 4.
(E-H) Membrane capacitance measurements of *FCHSD2*-KO cells.
(E) Average traces of 13 Control and 12 *FCHSD2*-KO cells. Control in black, *FCHSD2*-KO in red, SEM in grey.
(F) Cumulative increase in exocytosis normalised to size of cells. The data are presented as average at each pulse ± SEM and overlapping data points.
(G) Kinetics of exocytosis analysed through the increment in membrane capacitance at each pulse. Data normalised to cell size.
(H) Comparison of the size of the cells prior stimulation of exocytosis.

**Figure S7. Deletion efficiency of variant region by CRIPSR-Cas9 genome editing**
(A) Representative data of SYBR$^{TM}$ Green qPCR analysis on wildtype and dVR genomic DNAs. Note that the rate of DNA inversion detected by primer 1+3 was about 34% of remaining wildtype genomic DNA (primer 1+2) in dVR cells, roughly represent 5.4% of wildtype genomic DNA in wildtype cells.
(B) Deletion efficiency. The deletion efficiency rate (%) was determined by: 1. calculating the remaining wildtype allele (primer 1+2) in dVR genomic DNA and compared with wildtype allele in SHAM control and 2. Taking away the rate of inversion (primer 1+3) in dVR cells. Data are mean ± SEM. *, $P < 0.05$; **, $P < 0.01$; ***, $P < 0.005$. N = 1.

**Supplementary Tables**

Table S1 eQTL analysis of *FCHSD2* and STAR10 gene expression in human islet samples

| Gene | Variant ID | Position (hg19) | Ref allele | Alt allele | MAF | Risk allele | Nominal P value | rs140130268 R2 (EUR) |
|---|---|---|---|---|---|---|---|---|
| *FCHSD2* | rs11603334 | Chr11:72432985 | G | A | 0.144 | G | 0.0127082 | 0.8934 |
| *FCHSD2* | rs1552224 | Chr11:72433098 | A | C | 0.144 | A | 0.0127082 | 0.8934 |
| *FCHSD2* | rs75896506 | Chr11:72429829 | G | A | 0.144 | G | 0.0127082 | 0.8934 |
| *STARD10* | rs11603334 | Chr11:72432985 | G | A | 0.144 | G | $2.98 \times 10^{-4}$ | 0.8934 |
| *STARD10* | rs1552224 | Chr11:72433098 | A | C | 0.144 | A | $2.98 \times 10^{-4}$ | 0.8934 |
| *STARD10* | rs75896506 | Chr11:72429829 | G | A | 0.144 | G | $2.98 \times 10^{-4}$ | 0.8934 |

Table S2 Transcription factor binding affinity at genetic variants

| **variant** | **gene** | **Score** | |
|---|---|---|---|
| | | Risk allele | Protective allele |
| rs79430446 | GATA3 | 4.6395 | 6.22367 |
| | GATA5 | 4.36019 | 8.01532 |
| | GATA6 | 6.53008 | 8.17938 |
| | MAFK | 6.15646 | - |
| | NRL | 5.70744 | - |
| | MAFF | 7.22087 | - |
| | TBP | 6.96517 | - |
| | GSC | - | 8.9786 |
| | GSC2 | - | 8.55269 |
| | BARX1 | - | 6.55349 |
| | BSX | - | 5.48796 |
| | EVX1 | - | 6.53802 |
| | FOXP3 | - | 5.80405 |
| | OTX2 | - | 9.73373 |
| | OTX1 | - | 8.01065 |
| | HOXB2 | - | 5.44904 |
| | HOXB3 | - | 4.68114 |
| | MEF2C | - | 8.10057 |
| | MEF2A | - | 7.64429 |
| | RHOXF1 | - | 6.92681 |
| | VENTX | - | 6.05976 |
| | POU3F4 | 5.81505 | 10.7694 |
| | POU5F1B | 5.84768 | 11.6522 |
| | POU5F1 | - | 8.37144 |
| | POU2F1 | - | 9.09201 |
| | POU3F1 | - | 7.69322 |
| | POU3F2 | - | 7.49311 |
| | POU3F3 | - | 7.84807 |
| | RFX5 | 7.526 | - |

| | | | |
|---|---|---|---|
| rs140735484 | SP1 | 7.6665 | 10.0444 |
| | HIC2 | - | 5.55379 |
| | SP3 | - | 5.90913 |
| | CDX2 | 5.8768 | - |
| rs7103836 | HLF | 7.50884 | - |
| | GATA6 | 5.78747 | - |
| | SOX10 | 10.8334 | 5.84552 |
| | THAP1 | 6.55426 | - |
| | SOX15 | 6.50967 | - |
| | RUNX3 | 5.15967 | - |
| | SOX13 | 5.39057 | - |
| | FOXP3 | 5.92425 | - |
| | BARHL2 | 7.58065 | - |
| rs613937 | GSC2 | 6.72964 | - |
| | OTX1 | 7.62999 | - |
| | NKX3-2 | 5.9364 | - |
| | EBF1 | 4.59291 | 9.45164 |
| | RFX5 | - | 5.60305 |

Table S3 Oligo sequence for EMSA assay

| | | |
|---|---|---|
| rs140130268 | Risk allele | TATTTGTGGTTTGTTTGTTTGTTTTTCCT |
| | Protective allele | TATTTGTGGTTTGTTTGTTTTTCCT |
| rs79430446 | Risk allele | GTATAAGATGAGCATGGAAC |
| | Protective allele | GTATAAGATTAGCATGGAAC |
| rs140735484 | Risk allele | AAATCCCCCAGTCCCAGGCA |
| | Protective allele | AAATCCCCC-GTCCCAGGCA |
| rs7103836 | Risk allele | TCTGCCACAGAGACATAACA |
| | Protective allele | TCTGCCACACAGACATAACA |
| rs613937 | Risk allele | CAGCCTCCTAAGCGGCCACA |
| | Protective allele | CAGCCTCCTGAGCGGCCACA |

Table S4 gRNA sequences for CRISPR-Cas9 genome editing

| | |
|---|---|
| VR_KO1 | GACCCCTGTGAGCTCCTCGT |
| VR_KO2 | AGCGACCACCAGCTAGGTTT |
| CBS1 | GCTTGGGTGGGGGTGCAGCC |
| CBS2 | GGGCAGTCAAGGGCACAGGA |
| CBS5 | TGCAGAAGAATGGTCACTAG |
| CBS7 | AGTCTCGGGATCGACACGTG |
| CBS8 | CGGCCACAACCACTAGGGGG |
| R2_KO1 | GTGTCCTTCACAGTGGCGAT |
| R2_KO2 | ACTCCACCTTGGTATTACAC |
| STARD10-1 | AGAGGCCGCCAGCTTCTCCA |
| STARD10-2 | AGTCTTGGTCATCGGGCACC |
| FCHSD2-1 | GCATCATGCAGCCGCCGCCG |
| FCHSD2-2 | GCCCTTACCTTCCTCGGCGG |

Table S5 Primer set for genomic DNA PCR amplification

| | |
|---|---|
| VRdel_F1 | CCATCTCCCCGACTCAGCCCAG |
| VRdel_R1 | GGGAGATCCGATTTTGAGTCCCTGC |
| VRdel_F2 | CGACTCAGCCCAGTCTCCTCC |
| CBS1&2_F | GCAGCCGTGGCCAACACACACACTTCC |
| CBS1&2_R1 | CTCGTGGTGGGGTGCTTGCTGAGG |
| CBS1&2_R2 | GGAGCCCAGAGATGCTGAGAAACTTGC |
| CBS5_F | CCACTTCCAACCCCAGAGAC |
| CBS5_R1 | CATACTCAGGGGGCCTTGTG |
| CBS5_R2 | CCTTGTGGGGAGGGTCTGGGAG |
| CBS7&8_F | CACCCCTGGATCTCATTTGATCCTCC |
| CBS7&8_R1 | GCAGTCCTTGAATCCTGATCCTTCCCTGG |
| CBS7&8_R2 | GGTCAGAAGGACGATGCCGAGCGC |
| R2_F1 | GTCAGAAGGCTGAGGCAAGAGGATGG |
| R2_F2 | CAGTGAGCTGAGATGTTGCCACAGC |
| R2_R | CCACTTTGGTGCCATGTGTGGCCTGG |
| STARD10_CRISPR_F | CACCCCAGCCCTGCTATAGGTCAGG |
| STARD10_CRISPR_R | GCCCCCAGCGCACTGATTCCCGTCC |
| STARD10_CRISPR_R2 | GCACTGATTCCCGTCCCCAAACCG |
| FCHSD2_CRISPR_F1 | CTCCCTCGTCTCCTCACACTCG |
| FCHSD2_CRISPR_F2 | TGCCGCCCGCTGGCCTGCTCC |
| FCHSD2_CRISPR_R | CCCAAGACGAGGGCGGTCACG |

Table S6 SYBR™ Green qPCR primer for assessment of VR deletion

| | |
|---|---|
| dVR_1 | CCTTCTGGGCTCCCACACAATGC |
| dVR_2 | CTGCCCCAAATGTTCAACACGC |
| dVR_3 | TGTTCAACACGCACTCATTCTTCACC |
| dVR_4 | GTTGCTGAATCCCCCAAGCTTCAG |

Table S7 Nested PCR primer set for 4C assay at VR region

| | |
|---|---|
| 4C_F1 | GGTATAGCCTGCTACTCAAAAGGTCCCTGG |
| 4C_R1 | TACCTCCTGCACTGAGATTCTCCATGAAGC |
| 4C_F2-XhoI | AATTTCTCGAGCTTCTTGTGCCAGGAAGTAAGC |
| 4C_R2-NotI | ATGTGCGGCCGCGGACATCTCCCCATTTCCAAGG |

Table S8 Taqman™ primer/probe for 3C assay at VR region and R13 region

| | |
|---|---|
| **VR Probe** | AGATGTCCTAAAGTGCTCATTGGGGGCATA |
| Constant primer | CAGTGAAAGGAGGAGCCCACC |
| NcoI Fragment | |
| -13 | ACTTCTGTGAGCTCCCTGAGG |
| -12 | ACCTTGTCCGCTCTCAGTCC |
| -11 | CTCTCAGAGCCTGTCTGAACATAGC |
| 6 | CCCTTGGGGCTCTGTAGAGG |
| 7 | CCAAACCCACACCTGGAACAGG |
| 8 | CCAGCTCCACCGCTCCAAGG |
| 9 | ACAGGCGTGAGCCATCATGC |
| **R13 probe** | CCAGGCCTGGCCCTGTGCTGGCTCCTGAGG |
| Constant primer | ACTTCTGTGAGCTCCCTGAGG |

| NcoI Fragment | |
|---|---|
| 12 | CCCAACCTTTTTGGCACCAGG |
| 13 | CCGTGATGTCATCACCCTCC |
| 14 | CCTCCTGCACTGAGATTCTCC |
| 15 | GCAGCTTATCTCAGATTGAGCCC |
| 17 | CCTGGGTCCCTAGGACTTTGG |
| 18 | CTGGCAGAGGTGGTTTGAGC |
| 19 | CGGAGCCTCCGCGGAGGACC |
| 20 | CCAGAACACCAGGGACTCACG |
| 21 | ACTCCCCAGCCAGGTGAAGC |
| 23 | CAAGCGTGAGCCACTGCACC |

Table S9 Primer set for SYBR™ Green qPCR assay at CTCF binding site (CBS)

| | |
|---|---|
| CBS1_F | AAAGTCACCCAGAATCCCCC |
| CBS1_R | CAGGGCTTGTCAGTCAGGAC |
| CBS2_F | ATCAGCCTCCAGGAAGACCA |
| CBS2_R | GGTGTCCCCCTCTGATACCT |
| CBS3_F | CTCAGCCACCACATGACCTT |
| CBS3_R | GGGAGTCCCATCACAGTGTC |
| CBS4_F | CCAGTCAGGGTCCATGTTGG |
| CBS4_R | GGTCTTCAAAGCCCTGTGGT |
| CBS5_F | CCACAAGCTGATGGGGTTGA |
| CBS5_R | ACCTGGAGGGAAGCTCAGAT |
| CBS6_F | CACCAAATCTCCCTCACCCC |
| CBS6_R | AATCTCCTTCACAGACGCCC |
| Neg_F | AAAGGCCACAAACTCCCCAT |
| Neg_R | ATTTGGCAGAGCTGAGCGTT |
| CBS7_F | CAGAACGATGACTGGACCGT |
| CBS7_R | GGCCGCTGTAAACACCAAAG |
| CBS8_F | CAGAACGATGACTGGACCGT |
| CBS8_R | GGCCGCTGTAAACACCAAAG |

Table S10 Primer set for DNA cloning of active enhancer region

| | |
|---|---|
| R2_F | ACTGAGCTAGCGCAGTGAGCTGAGATGTTGC |
| R2_R | ACTGACTCGAGATCTGGCAACTCCACTTTGG |
| R4_F | ACTGAGCTAGCGAGTGGTCTGTTGCAGTCAGC |
| R4_R | ACTGACTCGAGGGTTTCACCACATTGGCCAGG |
| R5_F | ACTGAGCTAGCGGACAGAGTAACCTTAAGACACAGG |
| R5_R | ACTGACTCGAGCCAGAGAGGTGATGAGTCTTGAGG |
| R6_F | ACTGAGCTAGCTGGAAGGACACAGAGCACAG |
| R6_R | ACTGACTCGAGTCTCTGCCTCCCTTTCTCAG |
| R7_F | ACTGAGCTAGCGCAGGTGAAGAAACTGAGGC |
| R7_R | ACTGACTCGAGAACGTACTCAAGCCCAGCAT |
| R8_F | ACTGAGCTAGCCCTGCTCCTTACAGCCTCAC |
| R8_R | ACTGACTCGAGCCCTGTTCTTTGCTGTCCTC |

Table S11 SYBR™ Green qPCR primer for assessment of R2 deletion

| | |
|---|---|
| dR2_1 | GCTGGCTCGGCCTTGAGAGG |
| dR2_2 | GGCAAACACTTAGATCCGGCTCC |

| | |
|---|---|
| dR2_3 | TTCTCCTAATGTCCTGTCACAGTCCC |
| dR2_4 | ACTTTGGTGCCATGTGTGGCCTGG |

**ABBREVIATIONS**

T2D, Type 2 Diabetes

GWAS, Genome-wide association study

R, Regulatory region

SNP, Single-nucleotide polymorphism

CHIP-seq: Chromatin immunoprecipitation and DNA sequencing

ATAC-seq, Assay for Transposase-Accessible Chromatin using sequencing

3C, Chromosome Conformation Capture

4C, Circular Chromosome Conformation Capture

CTCF, CCCTC-bind factor

CBS, CTCF binding site

pcHi-C, Promoter Capture Hi-C

TAD, Topologically Associating Domain

STARD10, StAR-related lipid transfer protein 10

FCHSD2, FCH and double SH3 domains protein 2

CRISPR, Clustered Regularly Interspaced Short Palindromic Repeats

Cas9, Endonuclease from *Streptococcus pyogenes*

GSIS, Glucose-stimulated insulin secretion
