## Supplementary figures and images for "Chromatin 3D interaction analysis of the *STARD10* locus unveils *FCHSD2* as a new regulator of insulin secretion"

### Supplemental Figures

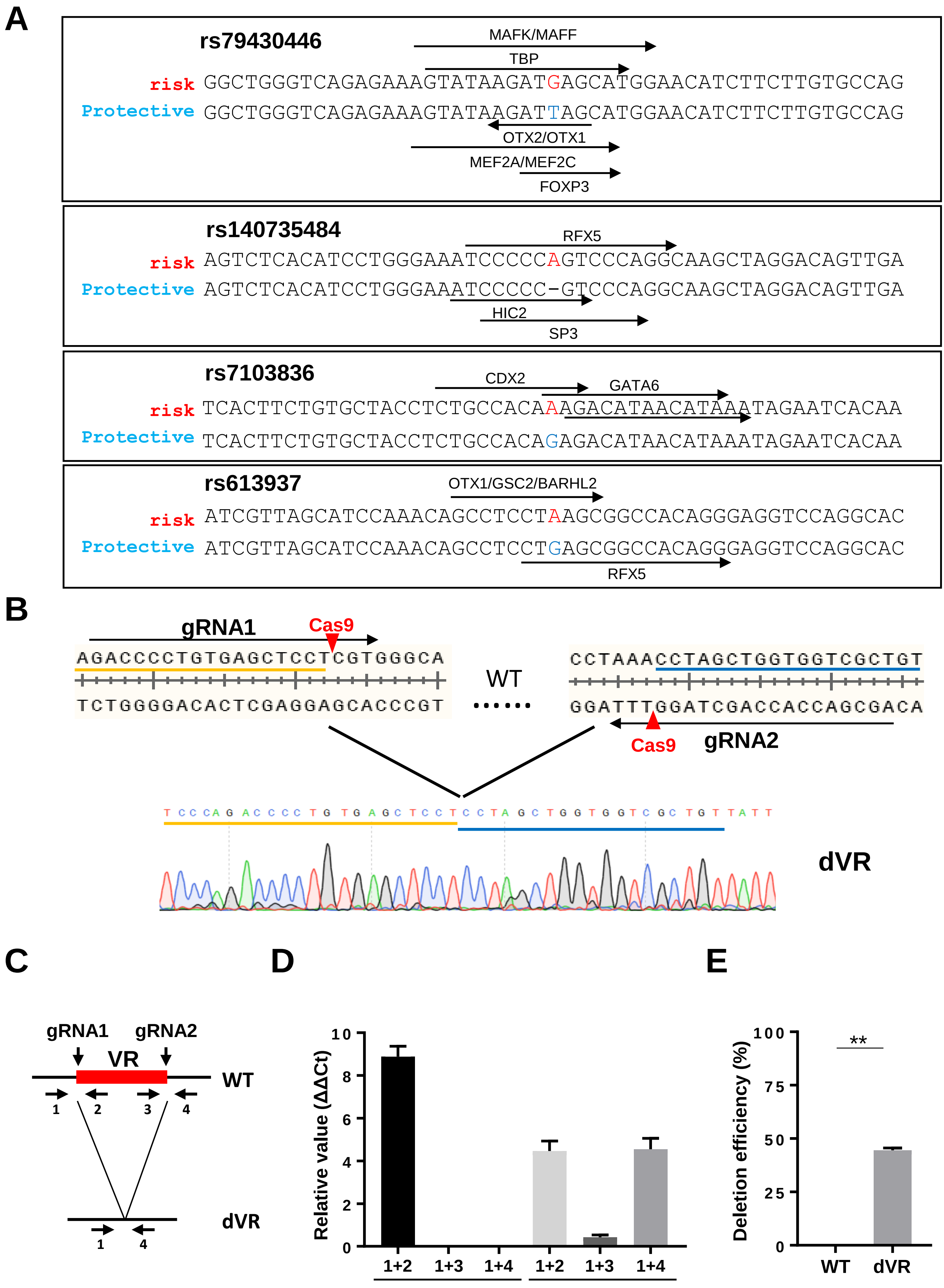

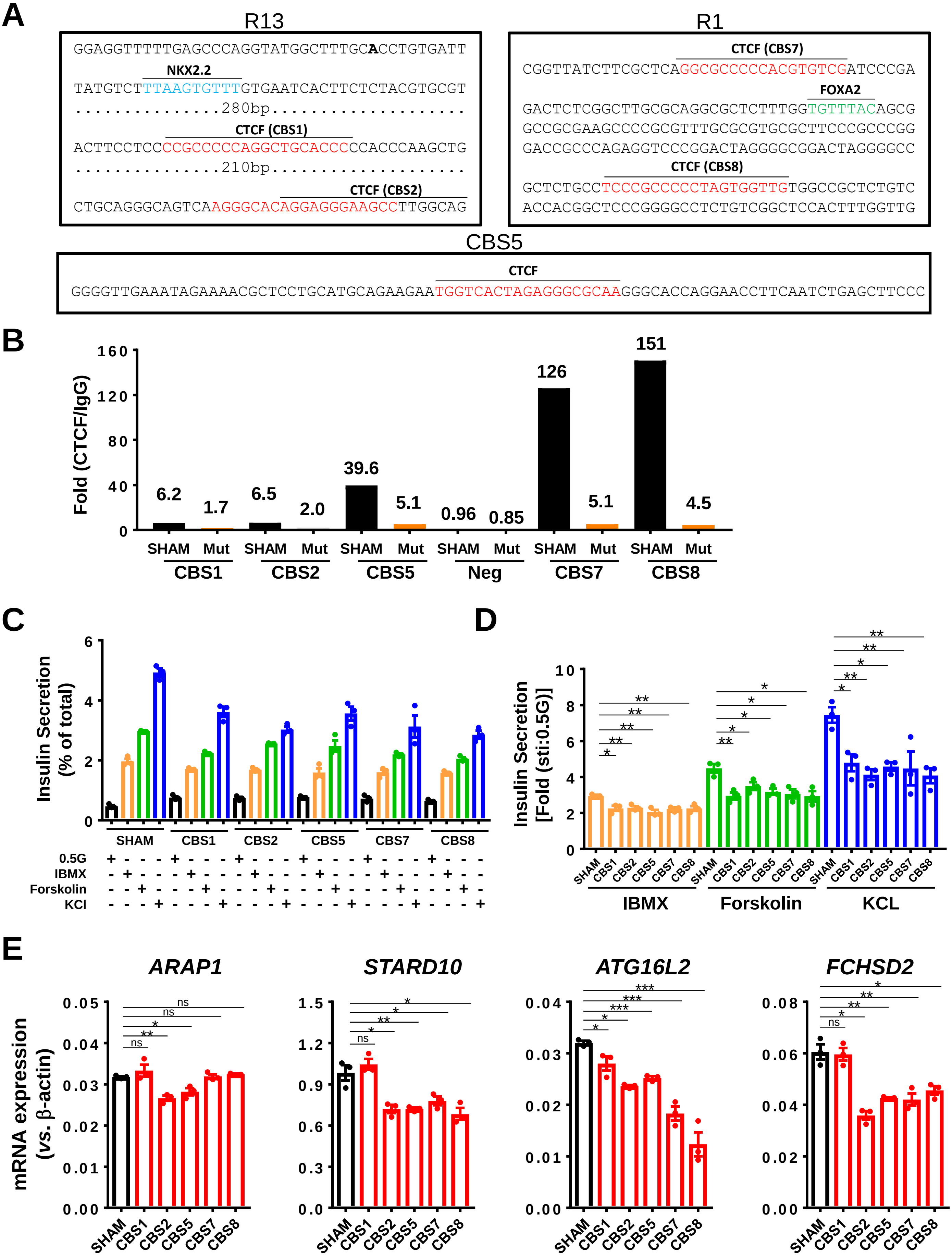

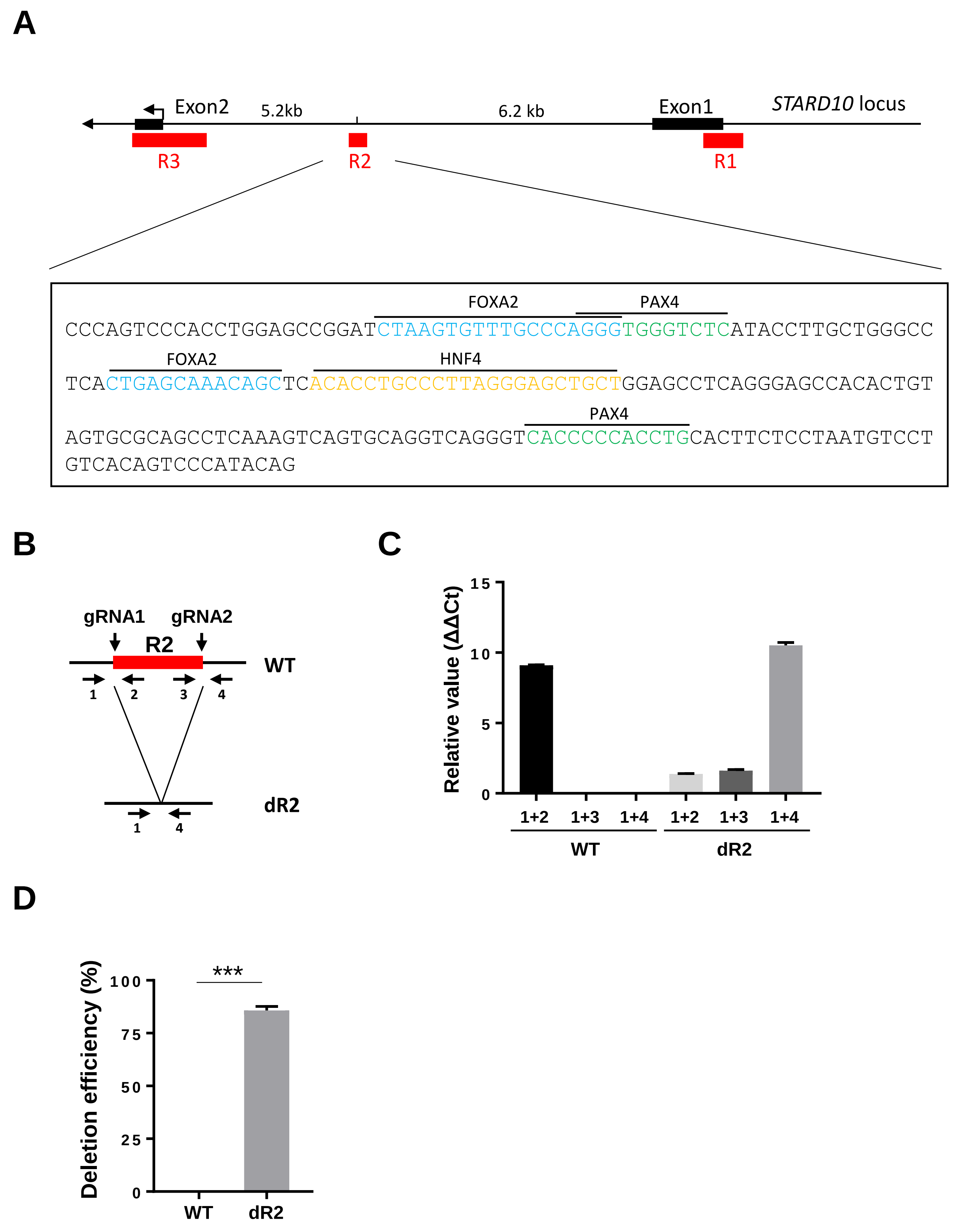

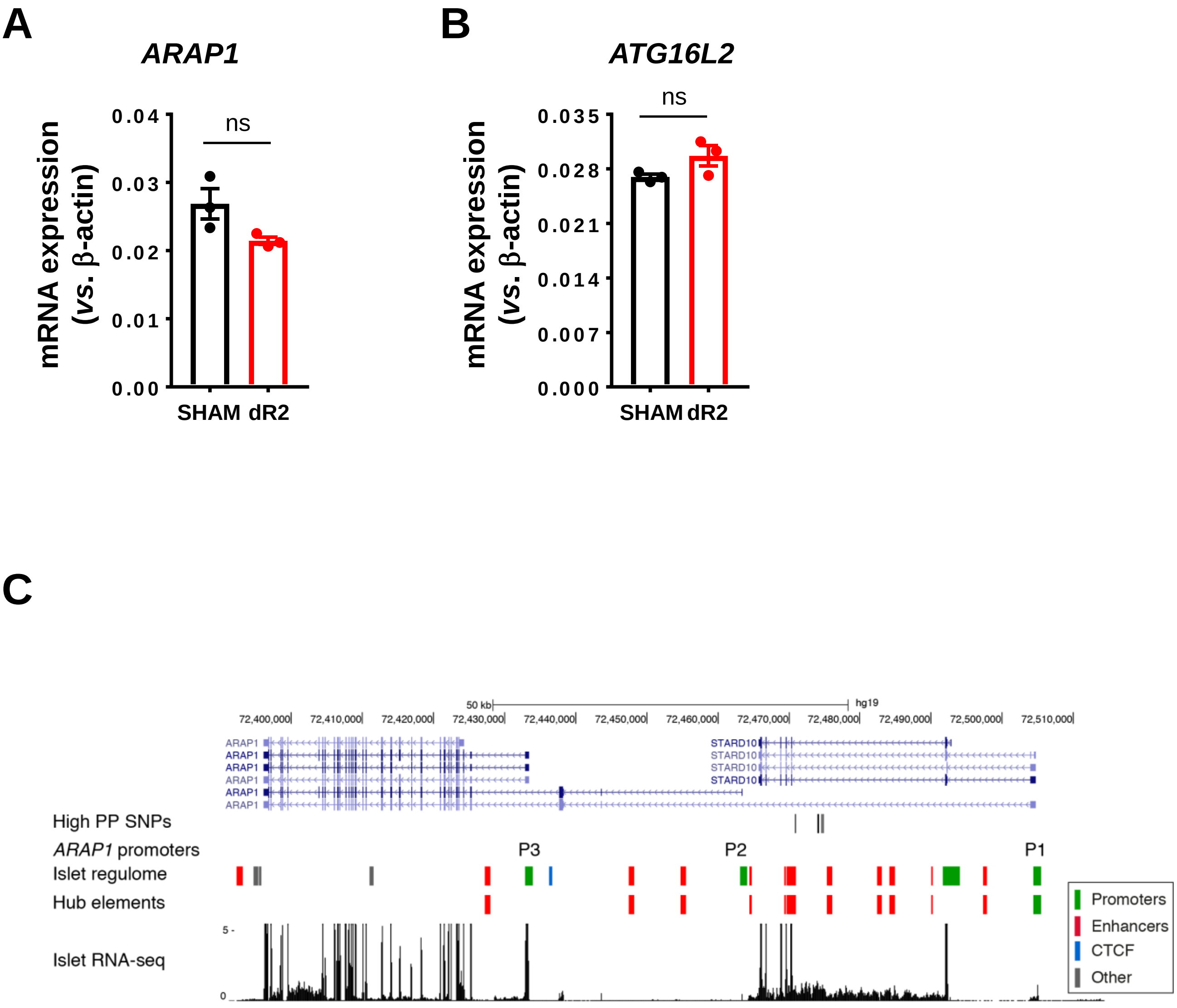

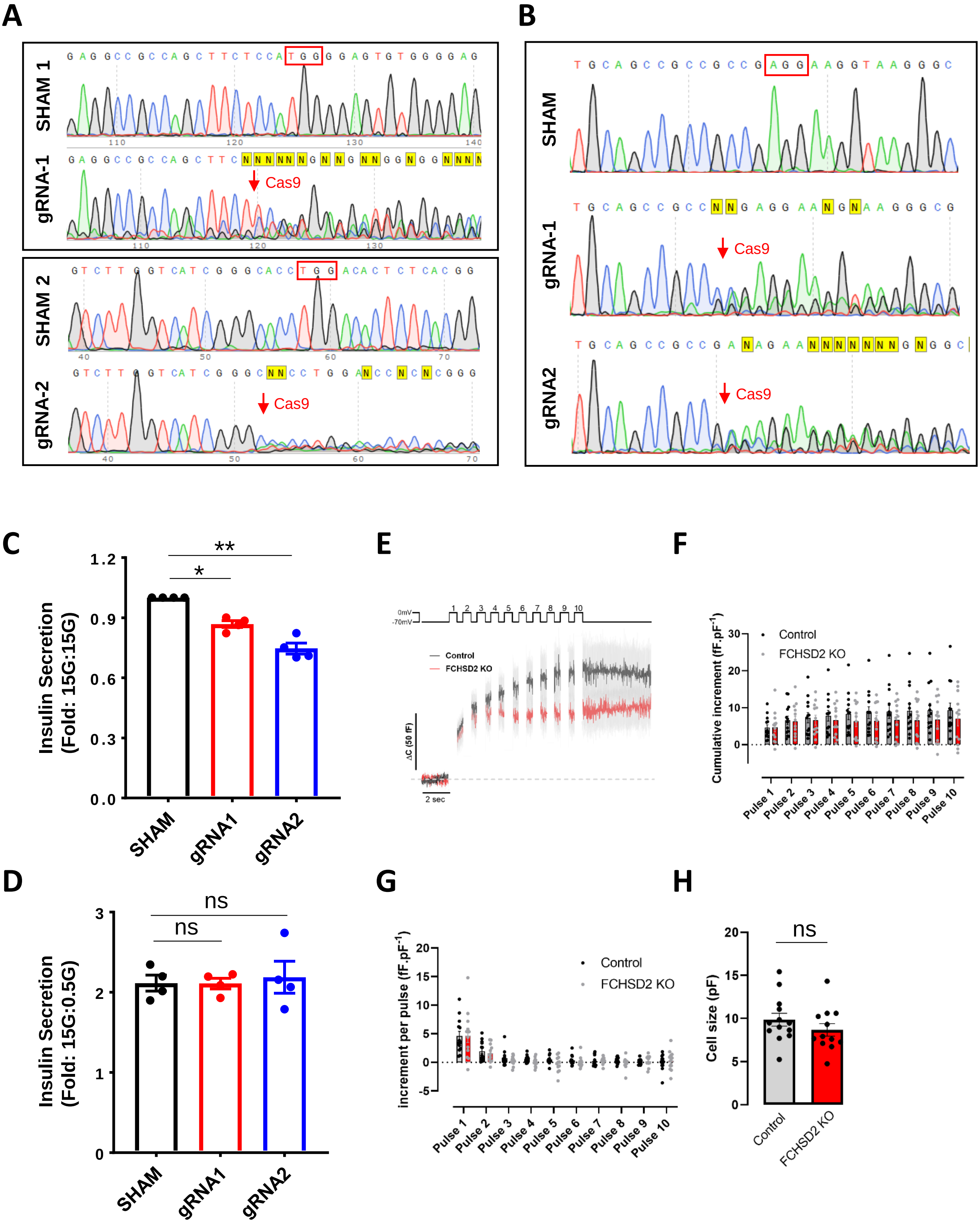

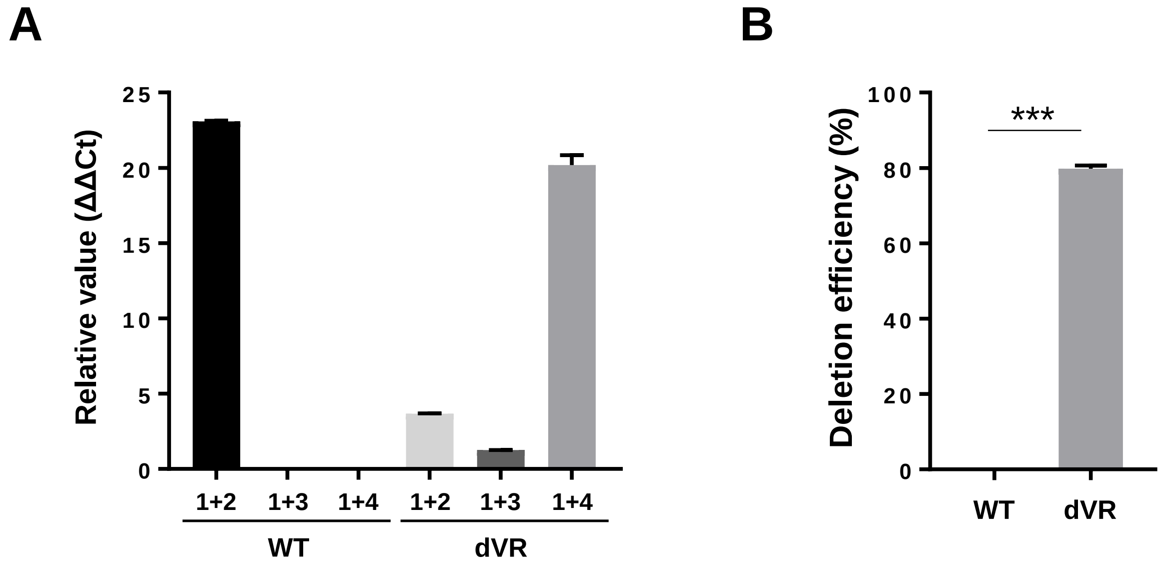
